# Functional Deorphanization of Tumor-reactive TCRs Uncovers a Landscape of Shared, Immune Dominant Antigens across Cancers

**DOI:** 10.64898/2026.09.28.755065

**Authors:** Orian Bricard, Razgar Seyed Rahmani, Frederik Peeters, Chloé Lorent, Thomas Van Brussel, Jieyi Xiong, Evy Vanderheyden, Rogier Schepers, Gino Philips, Eva Lion, Jiska van der Reest, Georg Halder, Ann Smeets, Michael Platten, Edward Wilhelm Green, Jeroen Dekervel, Diether Lambrechts

## Abstract

Single-cell sequencing has charted the clonal landscape of tumor-infiltrating T cells, but the antigens that drive these responses remain largely undefined, limiting rational immunotherapy design. Here, we introduce TWISTAR, a platform that systematically deorphanizes single-cell-derived TCRs against tumor transcriptomes. By directly validating TCR engagement, TWISTAR exclusively recovers functionally immunogenic antigens. Applying TWISTAR to melanoma, breast, and hepatocellular carcinoma patients responding to immune checkpoint blockade, we uncover that tumor-reactive TCRs recognize a broad repertoire of immunogenic targets beyond neoantigens, including peptides derived from unannotated coding sequences and novel transcripts. Many of these antigens are shared across patients and cancer types, revealing communal targets. Furthermore, we observe immune dominance, consisting of multiple clones converging on few immune dominant epitopes. Together, these findings reveal an unexpectedly broad landscape of tumor immunogenicity, expanding opportunities for the development of pan-cancer vaccines.

**HIGHLIGHTS:** • TWISTAR deorphanizes TCRs and recovers functionally immunogenic antigens

• Tumor-reactive TCRs recognize a broad repertoire of non-canonical tumor antigens

• Many non-canonical antigens are shared across patients and cancer types

• Diverse TCR clones converge on immune dominant epitopes in immunotherapy responders

**ETOC BLURB:** TWISTAR deorphanizes TCRs against autologous tumor transcriptomes, recovering functionally immunogenic antigens. Applied to immune checkpoint responders, it reveals shared non-canonical antigens across cancers and immune dominance, providing a roadmap for the development of pan-cancer vaccines.

## INTRODUCTION

Single-cell transcriptomics combined with T cell receptor (TCR) sequencing has revolutionized our ability to integrate transcriptional states of individual T cells with their TCR repertoires. By tracking which T cells clonally expand during immune checkpoint blockade (ICB), this has yielded key insights into the transcriptional signatures that underlie T cell activation and exhaustion^1–4^. Despite this progress, a critical gap still remains: the specific antigens recognized by these T cells are largely unknown. Deciphering this antigen landscape is essential to fully understand what drives anti-tumor immunity and develop more targeted, effective immunotherapies.

Some tumor-infiltrating T cells have been shown to recognize neoantigens, which arise from somatic mutations that give rise to mutant peptides specifically expressed by cancer cells. Neoantigens can be *in silico* predicted by leveraging sequencing data derived from the tumor. Recent clinical trials employing personalized mRNA vaccines targeting neoantigens have shown compelling results in various cancers such as melanoma, breast and pancreatic cancer^5–7^. Nevertheless, in a subset of patients, these prediction-based approaches fail to yield sufficient immunogenic neoantigens, rendering personalized vaccines ineffective. In such cases, characterizing non-mutated yet immunogenic tumor-associated antigens (TAAs) presents an attractive alternative. TAAs are typically classified based on their differential gene expression profile across tumors compared to non-malignant tissues, and they include antigens encoded by cancer testis genes (e.g., the MAGE family), tissue-restricted genes (e.g., tyrosinase), and genes that are overexpressed in tumors (e.g., HER2)^8^. Tumor antigens may also originate from alternative transcription start sites or aberrant splicing events, producing non-canonical transcripts with potentially high tumor specificity. Recent deep profiling of peptides presented by tumor HLA molecules using immunopeptidomics revealed a large array of putative antigens derived from non-canonical transcripts, so-called cryptic or dark antigens^9–11^. Although promising, the immunogenicity of these presented peptides and their relevance as therapeutic targets remains unproven.

Several approaches have been developed to deorphanize TCRs and identify their cognate antigens. TCR-antigen pairs are commonly identified using peptide-HLA multimers, small-scale peptide library screening, and *in silico* TCR-antigen prediction models^12–14^. While these methods are valuable to detect reactivity to known or predicted antigens, they typically capture only a limited fraction of T cell specificities, leaving substantial blind spots in our understanding of the full antigen landscape. Broader antigen discovery can be pursued using screening platforms based on large libraries, such as those representing the entire human proteome. Notable examples include TSCAN^15^ and TCR-MAP^16^. However, these libraries are constrained by current genome annotations, as they only include sequences that are known or predicted to be translated, but do not encompass patient-specific antigens not represented in reference databases. An alternative approach involves screening vast numbers of randomly generated peptides bound to HLA molecules through yeast display systems^17^. This method often yields sets of peptides sharing structural features, enabling the identification of critical amino acid residues and motifs essential for TCR recognition. However, these peptides rarely correspond to naturally occurring sequences, necessitating additional *in silico* research to identify potential matches between predicted motifs and the actual presented antigen. Collectively, these current platforms have captured only a narrow spectrum of TCR-antigen interactions, underscoring a pressing need for more comprehensive approaches.

Here, we developed TWISTAR (<u>T</u>ranscriptome-<u>WI</u>de <u>S</u>creens enabling <u>T</u> cell <u>A</u>ntigen <u>R</u>esearch), a novel TCR deorphanization platform that screens TCRs against a transcriptome library derived from autologous patient tumor RNA. Specifically, based on single-cell RNA and TCR profiling, we systematically selected tumor-reactive TCRs from T cells identified within 3 distinct clinical contexts: (1) tumor-reactive, tumor-infiltrating lymphocytes (TILs) derived from a melanoma patient with brain metastasis; (2) PD1-positive, clonally expanded TILs from an invasive breast carcinoma patient treated with ICB; and (3) dominant T cell clonotypes from a hepatocellular carcinoma (HCC) patient experiencing a complete pathological response following ICB. For 40 out of 61 TCRs selected, TWISTAR successfully identifies the cognate antigens, validating a total of 31 presented epitopes. Interestingly, several of these antigens originate from non-canonical transcripts that were frequently expressed in various other tumors. Moreover, several of these antigens were identified by multiple clonal T cells with different TCR sequences, revealing an unprecedented level of immune dominance driving anti-tumor immune responses.

## RESULTS

### Engineering TWISTAR: a High-Sensitivity Screening Platform for TCR Antigen Discovery

To identify the cognate antigen of any given TCR, we developed TWISTAR. It is comprised of two key steps: an initial screening and a subsequent hit identification step. To screen, GFP-reporter T cells engineered to express a TCR of interest are co-cultured with millions of 293T cells transduced with patient’s HLA alleles and an autologous tumor-derived lentiviral cDNA library. Each 293T cell acts as an antigen-presenting cell (APC) expressing a fragment of the tumor transcriptome and presenting potential antigenic peptides derived from it. When T cells recognize an APC presenting their cognate antigen, a cluster of activated GFP+ T cells is formed, which is detected via an automated microscopy platform (**Figure 1A**). In the hit identification step, APCs surrounded by GFP+ T cells are retrieved, using a purpose-built cell picker mounted on the microscopy platform, and re-seeded for amplification and validation through a secondary co-culture with reporter T cells. Subsequently, antigen-encoding sequences are retrieved by sequencing the tumor-derived transcriptomic fragment integrated in the APCs. Finally, the epitopes are predicted computationally for their ability to bind HLA and validated through peptide-pulsed recognition assays (**Figure 1B**).

**Figure 1.**
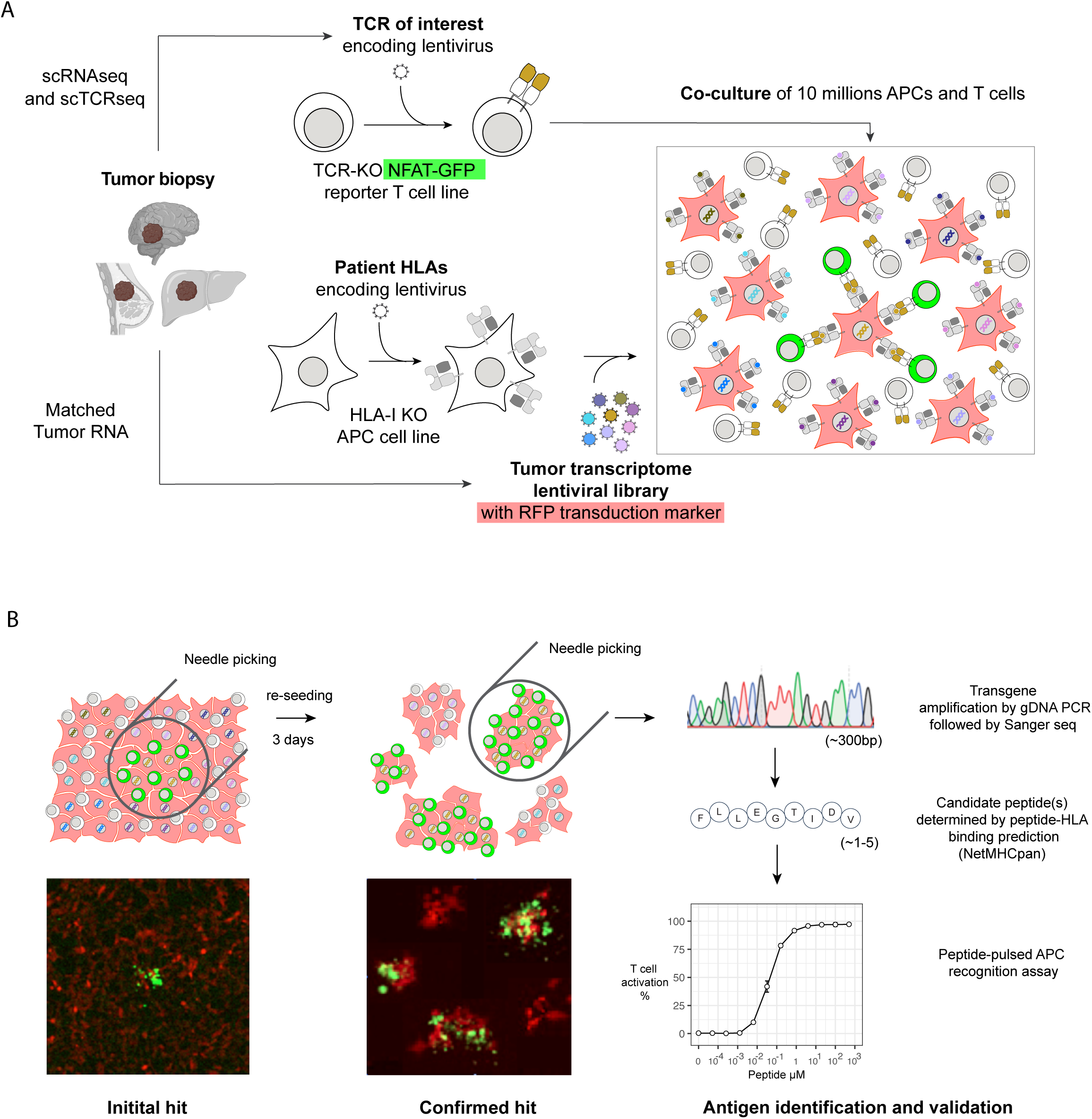
TWISTAR: an Autologous Tumor RNA-based TCR Deorphanization Platform. (A) Initial screening: tumor-derived RNA is converted into a lentiviral cDNA library (RFP-labeled) and transduced into HLA-I expressing APCs. Reporter T cells, engineered to express a TCR of interest and a NFAT-GFP activation reporter, are co-cultured with the APC library. Recognition of cognate antigen triggers GFP expression, generating detectable GFP+ T cell clusters (schematically shown as green clusters interacting with red APCs). (B) Hit identification and validation: APCs surrounded by GFP+ T cells are collected via automated cell picking, expanded and validated through a subsequent co-culture with reporter T cells. The integrated transcript fragment (∼300bp) is recovered by PCR and identified by Sanger sequencing. Candidate peptides (∼1-5) derived from the identified ORFs are selected based on their predicted ability to bind patient HLAs and validated in peptide pulse recognition assays to confirm TCR specificity.

The recognition of peptide-HLA complexes by TCRs requires that a protein is expressed at sufficiently high level to be processed into peptides, loaded onto HLA molecules, and presented for TCR engagement. At the transcript level, a threshold of at least 1 transcript per million (TPM) is commonly accepted as the minimum required for engagement^18, 19^. Consequently, a cDNA-based antigen discovery platform must interrogate millions of antigen-presenting cells (APCs), each expressing a single tumor transcript fragment, to adequately represent the entire tumor transcriptome (see **Methods**). Screening at this scale presents considerable challenges in both signal sensitivity and specificity. To address this, we turned to Jurkat reporter T cells that express GFP upon TCR-mediated activation^20^. While these Jurkat T cells typically rely on stimulation by many cells to induce GFP expression, we observed that they can also be activated by a single APC. Specifically, upon co-culture with adherent 293T APCs and following TCR engagement, the activated Jurkat cells remain localized around the APC they recognize, forming distinct GFP+ clusters. These are easily distinguishable from the low-level, leaky GFP expression that occurs in ∼0.5% of Jurkat cells growing in suspension. Furthermore, because these Jurkat cells lack cytolytic activity, the recognized APCs remain viable, enabling the subsequent recovery and sequencing of the integrated cDNA fragment that encodes the antigen. In TWISTAR, we exploited this property to enable large-scale screening of millions of APCs transduced with tumor-derived cDNA libraries.

To enable efficient screening, several technical optimizations were implemented. First, while conducting proof-of-concept experiments using a high and low-affinity TCR specific for a mutant PIK3CA peptide (mutPIK3CA+)^21^, we observed the systematic formation of GFP+ clusters upon antigen recognition for the high, but not the low-affinity TCR (**Figure S1A**). This was linked to genetic drift in the Jurkat-76 NFAT-GFP reporter line^22^, leading to loss of adhesion molecules such as CD2 and CD226, which result in suboptimal signaling in case of the lower-affinity TCR^23^. Because Jurkat-E6.1 retains expression of these molecules, we generated a novel reporter cell line based on this parental line. We disrupted endogenous TCRα/β sequences using CRISPR/Cas9, integrated multiple copies of a NFAT-GFP reporter cassette, and stably expressed the TCR co-receptor CD8. Multiple clonal lines were derived and tested for their ability to induce GFP when transduced with the low-affinity TCR and cocultured with mutPIK3CA+ cells. One clone (referred to as Sirius Line 8 or SL8) showed the strongest propensity to form intense GFP+ clusters (**Figure S1B,C**) and was selected for subsequent TWISTAR screening experiments. Additionally, to optimize TCR expression into the reporter T cell line, we compared seven retroviral and lentiviral vectors. Amongst them, a lentiviral backbone containing an EF1α promoter and an intronic sequence encoded upstream of the TCR coding sequence (pWPXL), enhancing mRNA processing and stability, produced the highest TCR expression levels (**Figure S1D**).

Second, we considered 293T cells as suitable APCs, as they express the core components needed for antigen processing and presentation, such as genes involved in MHC class I presentation and proteasome degradation^24^. To prevent potential reactivity towards patient-irrelevant HLA alleles, we generated a 293T HLA-I null line, in which we re-expressed HLA alleles by lentiviral integration. Each individual HLA was transduced in an independent APC line to prevent inter-allelic competition for antigen presentation. To enhance antigen presentation ability, we explored whether overexpression of peptide-loading complex components increases surface HLA expression and T cell activation. Using lentiviral transduction, we stably overexpressed *B2M*, *TAP1*, *TAP2*, *TAPBP*, *CALR* and *PDIA3*. We observed that B2M, TAP1 and TAPBP enhanced HLA expression in 293T cells and subsequent T cell activation of Jurkat cells (**Figure S1E,F**). 293T APCs overexpressing B2M, TAP1, and TAPBP were therefore maintained in subsequent experiments.

Finally, for the generation of tumor-derived cDNA transcriptome lentiviral libraries, we reasoned that full-length cDNA cloning would bias the library, due to vector size constraints favoring short transcripts. We therefore implemented a cloning protocol starting from randomly fragmented cDNA, in which transcripts are fragmented to generate ∼300 bp inserts. These were cloned into a lentiviral expression cassette and compared to the original tumor transcriptome assessed by high-coverage RNAseq. We retrieved a similar representation for annotated and unannotated genomic loci in the corresponding TWISTAR library (Pearson correlation >0.99) (**Figure S1G**).

### TWISTAR allows Deorphanization of Tumor Cell Line Reactive TCRs

To evaluate TWISTAR’s ability to identify cognate antigens from tumor-specific TCRs, we analyzed 30 TCRs triggering lytic activity when expressed in primary T cells co-incubated with the BT21 melanoma cell line. All TCRs were identified by single-cell profiling of tumor-infiltrating lymphocytes (TILs) from the primary melanoma-derived brain metastasis of which the BT21 cell line was derived^25^. As a negative control, we tested 20 TCRs derived from the same tumor, but with no detectable reactivity towards BT21. All 50 TCRs were identified as abundant T cell clonotypes infiltrating the metastatic lesion (**Figure 2A**) and were characterized by an activated phenotype with exhaustion based on expression of effector (*IFNG*), cytotoxic (*PRF1, GZMB*) and immune-checkpoint (e.g. CTLA4, *PDCD1, TIGIT)* molecules, and the chemoattractant *CXCL13* (**Figure 2B**). Overall, these 50 TCRs provide a benchmark to assess TWISTAR’s sensitivity and specificity to identify cognate tumor antigens in settings where tumor reactivity has been established but the target antigens remain unknown.

**Figure 2.**
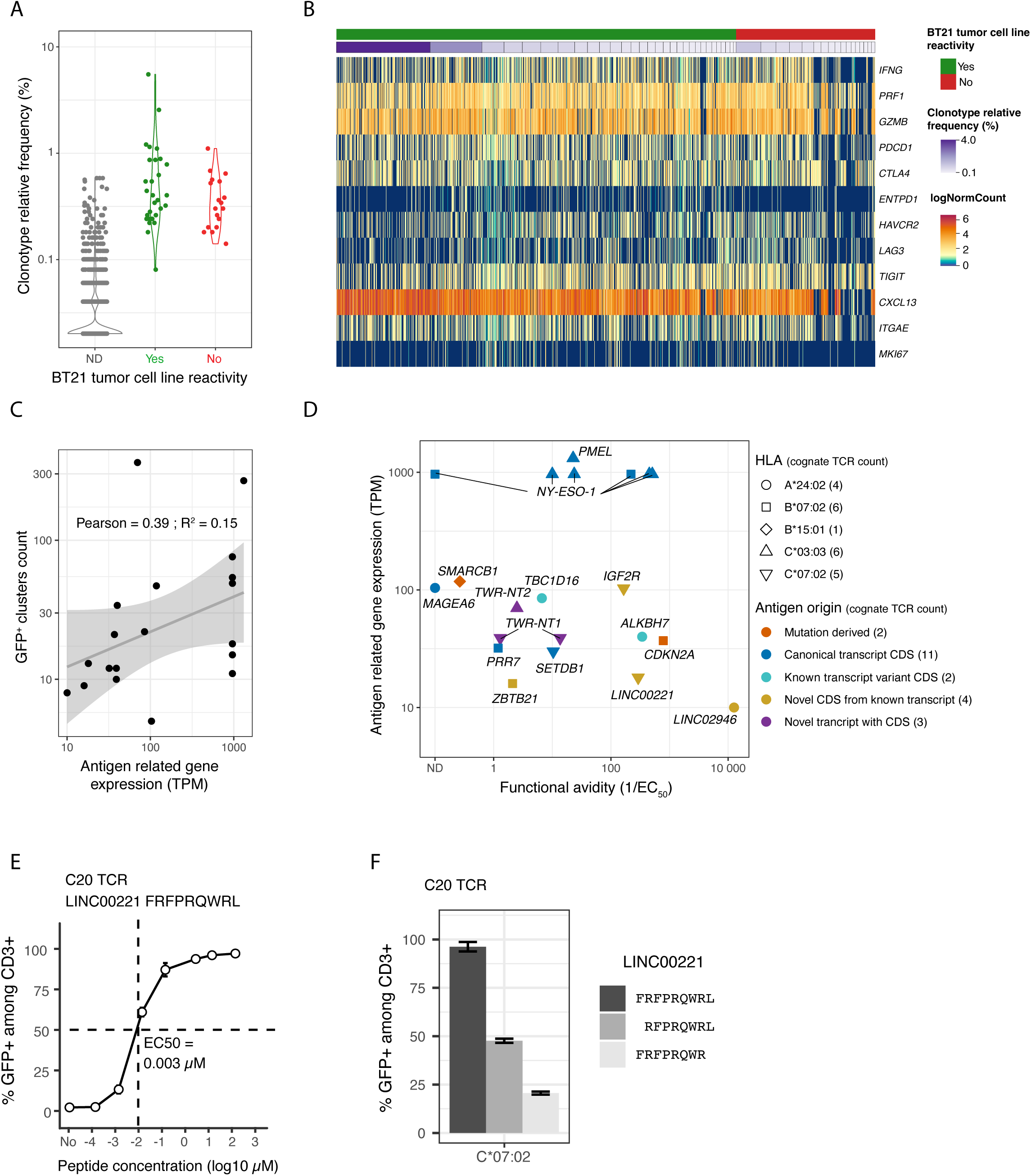
Deorphanization of BT21 Tumor Cell Line Reactive TCRs. (A) Frequencies of T cell clonotypes from BT21 tumor-infiltrating lymphocytes (TILs) based on reactivity of their TCRs towards the BT21 tumor cell line. (B) Single-cell gene expression heatmap with rows representing individual genes and columns representing individual cells, grouped according to reactivity of their associated TCRs towards the BT21 tumor cell line and their clonotype relative frequency. (C) Correlation between antigen-related gene expression and the number of GFP+ clusters identified in the initial screening by TWISTAR. Each point reflects expression levels (in transcripts per million or TPM) on the X-axis and the number of GFP+ reporter clusters detected. The solid line indicates the linear regression fit, with the shaded area representing the 95% confidence interval. Pearson correlation coefficient (r) and coefficient of determination (R²) are indicated. (D) Characteristics of antigens recognized by deorphanized TCRs from BT21 TILs (n=22). Each dot represents the cognate antigen of one deorphanized TCR. The graph depicts functional avidity defined as the inverse of the half-maximal effective concentration (EC_50_, µM) relative to antigen-related gene expression determined by bulk RNAseq of the BT21 cell line. HLA restriction of each TCR-antigen pair is indicated by a symbol, while the type of transcript from which the antigen is derived is indicated in color. Number of cognate TCRs associated with each subgroup is indicated in parentheses. (E,F) Validation and characterization of the LINC00221-derived epitope. HLA-C*07:02-expressing cells were pulsed with the indicated peptides at varying concentrations (E) or at 50 μM (F) and co-incubated with reporter T cells transduced with the TCR recognizing the LINC00221-derived epitope (C20). Peptide titration assays were used to determine EC_50_ values (E), while peptide truncation analysis defined the minimal epitope length supporting maximal TCR recognition (F). GFP expression was measured using flow cytometry. Measurements were performed in duplicate. Error bars represent 95% confidence intervals. The number of cognate TCRs associated with each subgroup is indicated in parentheses.

Out of the 20 non-reactive TCRs screened against the custom BT21-derived cDNA library, we observed a small number of weak GFP+ clusters, which after picking and re-seeding, did not result in a population of APCs recognized by the corresponding TCR (**Supplementary Table 1**). This indicates that TWISTAR has a low likelihood of identifying false-positive tumor antigens. In contrast, when screening the 30 tumor-reactive TCRs, numerous GFP+ clusters were detected, ranging from 1 to 362 per TCR investigated on ∼10 million seeded APCs (**Supplementary Table 1**). After picking and re-seeding, most clusters gave rise to APCs capable of triggering strong GFP signals in T cells expressing the corresponding TCRs. Sequencing of the cDNA integrated in these APCs resulted in the identification of antigen encoding sequences for 22 of the 30 TCRs investigated (73%). Such a high TCR-antigen deorphanization rate has to the best of our knowledge not previously been reported.

The tumor antigens we identified originated from 15 genes expressed in BT21 at levels ranging between 10 to 964 TPM. We observed a moderate positive correlation between the number of GFP+ clusters and expression of the gene encoding the cognate antigen, suggesting that screening sensitivity is at least partially determined by the extent to which transcripts are represented in the antigen library (**Figure 2C**). Each of the 5 HLA-I alleles expressed in BT21 (which has a homozygous HLA-A) presented a peptide recognized by at least one of the TCRs (**Figure 2D**). Next, NetMHCpan was used to predict which peptides within the ±300 nucleotide cDNA inserts were presented by each corresponding HLA allele. When multiple candidate peptides were predicted, we subcloned the cDNA insert into smaller fragments and tested which cDNA fragment gave rise to TCR recognition. Finally, all peptides predicted by NetMHCpan were confirmed using an orthogonal approach, whereby for each predicted peptide varying concentrations of synthetic peptides were pulsed onto 293T cells expressing a single HLA allele and tested for recognition by T cells transduced with the corresponding TCR (**Figure S2A**). From these assays, we determined the half-maximal effective concentration (EC_50_), representing the peptide concentration required to elicit 50% of maximal T cell activation. The EC_50_ concentrations ranged from 0.08 nM to 3.8 μM, reflecting a broad spectrum of functional avidities. We also confirmed the minimal peptide length eliciting the strongest T cell response by testing shorter peptides trimmed by one amino acid on either side of the predicted peptide (**Figure 2E,F and S2B**). Overall, this strategy reliably identified the exact epitopes for 19 out of 22 deorphanized TCRs (**Supplementary Table 1**).

Among the 22 deorphanized TCRs, only 2 recognized classical neoepitopes derived from somatic mutations in *CDKN2A* and *SMARCB1*. Of the remainder, 11 epitopes were derived from canonical protein coding sequences (CDS): MAGEA6, NY-ESO-1 (also referred to as CTAG1A/B), PMEL (Gp100), PRR7 and SETDB1. NY-ESO-1 appeared to be immune dominant, with 7 TCRs recognizing 4 different epitopes of this protein presented on two different HLA alleles. Two TCRs recognized epitopes derived from a transcript variant located within an annotated CDS (*e.g.,* an alternative upstream translation initiation start site in *ALKBH7* gives rise to the epitope). Four TCRs recognized epitopes derived from a novel CDS within previously described transcripts (e.g., the annotated lncRNA *LINC00221* contains an unexpected CDS giving rise to an epitope) and 3 TCRs recognized epitopes derived from newly identified transcripts (*e.g., TWISTAR-New Transcript 1* or *TWR-NT1*, is related to a set of novel transcripts overlapping with *ENSG00000302645* and *FOXR2* on chromosome X). Three out of 4 epitopes derived from NY-ESO-1 have previously been associated with TCR reactivity^26^. The 5 epitopes originating from SETDB1, PMEL, LINC00221, TBC1D16, and ALKBH7 have been reported in immunopeptidome databases^10, 26^, but based on TWISTAR could now be associated with specific TCR reactivity. The 2 epitopes derived from IGF2R and PRR7 are reported here for the first time, although peptides originating from the same coding frame have previously been reported in immunopeptidomic datasets. Finally, the remaining 4 epitopes represent previously unreported antigens (**Supplementary Table 1**). Together, these findings uncover an unexpectedly broad and mechanistically diverse immunogenic landscape, in which non-mutational antigens, derived from non-canonical transcripts and alternative translation products, feature prominently alongside neoantigens.

### TWISTAR Deorphanizes Patient-specific Mutation-derived Tumor Neoantigens

TWISTAR revealed that only 2 of the 22 deorphanized TCRs recognized neoantigens arising from somatic mutations. Both derived from tumor suppressor genes by single-base substitutions but were not annotated as hotspot mutations. Notably, in case of *SMARCB1*, the mutation introduced an amino-acid substitution within the central region of the presented peptide, generating a clear tumor-specific epitope recognized by the cognate TCR (**Figure 3A,B**). In contrast, the mutation in *CDKN2A* introduced a premature stop codon downstream of the antigenic peptide coding sequence. Consequently, the presented epitope corresponds to the wild-type sequence (**Figure 3C**). However, overexpression of the full-length wild-type *CDKN2A* coding sequence did not trigger TCR recognition, while expression of the mutated allele did (**Figure 3D**), suggesting that the mutation leads to production of a truncated protein that is differentially processed by the proteasome, thereby enabling presentation of a peptide that is otherwise not displayed on wild-type cells. Importantly, this mutation was identified by TWISTAR but cannot be identified by standard neoantigen prediction pipelines as they do not take altered proteasome processing into account.

**Figure 3.**
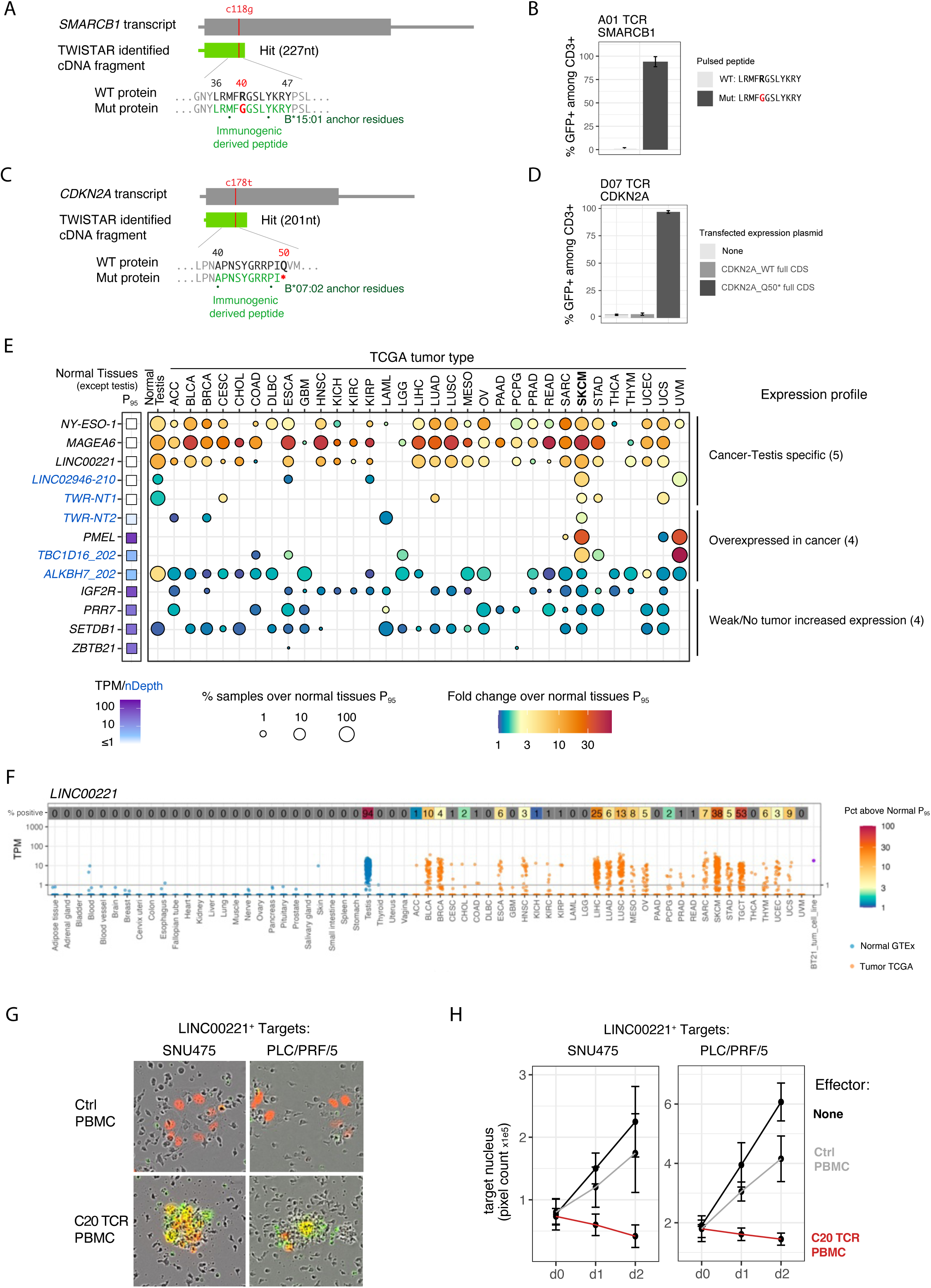
Mutation-derived and Tumor-associated Antigens of BT21 Tumor-infiltrating Lymphocytes. (A,B,C,D) T cell activation in response to *SMARCB1* and *CDKN2A* mutation-derived antigens. (A,C): Mapping of the presented epitope relative to the somatic mutation (transcripts with the base substitution in red and the cDNA insert of the APC in green). Epitopes derived from wild-type (WT) and mutant (Mut) proteins are indicated. The stop codon in the *CDKN2A* mutant is indicated by an asterisk. The recognized epitope is highlighted in green, with anchor residues indicated by dots. (B) HLA-B*15:01 cells were pulsed with *SMARCB1* WT or mutant peptides and co-incubated with reporter T cells transduced with the corresponding cognate TCRs (A01 and D07). (D) HLA-B*07:02 cells were transfected with plasmids encoding either the full-length WT *CDKN2A* or Mut *CDKN2A_Q50\** coding sequence. (B,D) The percentage of GFP+ cells among CD3+ T cells after a co-culture with TCR-transduced reporter T cells. Measurements were performed in duplicate. Error bars represent the 95% confidence interval. (E) Expression profile of antigen-related genes across normal tissues from the Genotype-Tissue Expression (GTEx) portal and tumors from The Cancer Genome Atlas (TCGA). In the left panel, the highest 95th percentile of expression across normal tissues (excluding testis) indicated by a color scale in squares is used as a threshold to assess increased expression in tumors. The minimal threshold value was fixed at 1 transcript per million (TPM). In the central panel, for each gene and cancer type, size represents the percentage of tumor samples with expression above the 95th percentile of normal tissues (P_95_), while color indicates the fold change relative to this threshold. Expression values are shown in TPM. Cancer types are abbreviated according to TCGA nomenclature. (F) *LINC00221* expression across normal tissues from GTEx (blue) and tumors from TCGA (orange). For each gene, the Y-axis shows expression in TPMs (log scale), and each dot represents an individual sample. The numbers and color above each tissue or cancer type indicate the percentage of samples with expression above the 95th percentile of normal tissues (P_95_), excluding testis. Cancer types are abbreviated according to TCGA nomenclature. LIHC stands for the TCGA Liver Hepatocellular Carcinoma dataset. (G,H) Functional validation of the LINC00221-derived epitope. *LINC00221*-positive SNU475 and PLC/PRF/5 cells expressing RFP were co-cultured with healthy donor PBMCs expressing the corresponding TCR (C20) or with untransduced control PBMCs. (G) Representative Incucyte images showing tumor cell growth (red nucleus) and T cell-mediated cytotoxicity (caspase 3/8 green dye). (H) Longitudinal quantification of tumor burden based on target nucleus counts. Cultures consisting of PBMCs transduced with the TCR exhibited sustained suppression of tumor growth compared to control conditions. Measurements were performed in duplicate. Error bars represent the 95% confidence interval.

### TWISTAR Identifies Tumor-reactive TCRs Targeting Shared Tumor Antigens

Interestingly, TWISTAR deorphanized 9 TCRs with reactivity towards MAGEA6, NY-ESO-1 and PMEL. These are cancer testis antigens commonly associated with immunogenicity, especially in melanoma where 69%, 26% and 79% of tumors exhibit high expression levels for these genes (**Figure 3E**). We found that antigen-related gene expression for antigens not derived from classical cancer testis antigens was also high in melanoma and other cancer types in TCGA, suggesting they represent shared tumor antigens across cancer types (**Figures 3E,F** and **S3C,D**). For instance, we observed specific expression profiles for *LINC00221*, *LINC02946* and *TWR-NT1*, with minimal to no expression in normal tissue except for testis, but high expression in a considerable fraction of tumors across cancer types. *LINC02946-210* (for which the epitope was derived from a 102 bp ORF) was observed in >60% of skin and uveal melanomas, while *LINC00221* (derived from a 66 bp ORF) was upregulated in 35% of melanoma tumors, but also in 25% of HCC patients and several other cancer types (13% of lung and 10% of bladder cancer patients; **Figure 3F**). Expression of the cognate TCR in primary T cells followed by co-culture with LINC00221^+^ HCC tumor cell lines (SNU475 and PLC/PRF/5) resulted in tumor cell killing and sustained growth suppression (**Figure 3G,H**), confirming LINC00221 as an immunogenically relevant target in cell lines derived from other cancers. Notably, LINC00221 knock down in HCC has been shown to reduce tumor growth, migration and invasion, suggesting that it acts as an oncogenic driver^27^. Peptide pulse experiments revealed that EC_50_ values for both lncRNA-related epitopes were low (0.08 and 3 nM), consistent with a permissive central tolerance preserving a high-affinity TCR repertoire.

We also identified a second set of genes (e.g., *TWR-NT2*, *TBC1D16-202*, and *ALKBH7-202*) that exhibited baseline expression in normal tissues, but substantial upregulation in tumors, suggesting these also represent tumor-associated antigens (**Figures 3E and S3D**). Intriguingly, their TCR avidities ranged considerably, with TCR avidity for the ALKBH7-derived epitope being similar to that of the *CDKN2A* mutation, challenging the prevailing notion that tumor-associated antigens are strictly recognized by low-affinity TCRs^28^. Overall, these findings reveal that most BT21 tumor antigens recognized by tumor-infiltrating T cells are derived from transcripts specifically upregulated in tumors and shared across patients and cancer types.

While most antigens followed the expected pattern of linear peptide presentation, MAGEA6 presented an exception. NetMHCpan predicted several peptides within the antigen encoding sequence of MAGEA6. When testing smaller fragments of the original cDNA insert, we found that none of the predicted peptides was compatible with recognition of these sequences. Indeed, the smallest insert that triggered recognition consisted of 207 nucleotides; much longer than expected for a linear HLA-displayed peptide. Both ends of the encoded 69-mer polypeptide contained amino acids matching the binding motif of HLA-A*24:02, suggesting that the recognition involves peptide splicing of the 69-amino acid polypeptide. The exact sequence of this epitope, however, remains to be determined. Since MAGEA3, MAGEA6 and MAGEA12 are highly similar in protein sequence, a TCR can recognize peptides derived from all 3 proteins. Therefore, we also tested corresponding sequences from MAGEA3 and MAGEA12 against our TCR, identifying reactivity against MAGEA3, but not MAGEA12^29^ (**Figure S3A,B**). This is noteworthy, given that infusion of anti-MAGEA3 TCR-T cells in the NCT01273181 clinical trial, which led to the death of two patients, has been associated with cross-reactivity towards MAGEA12 expressed in the brain^30^.

### Complex Regulatory Events Underlie Tumor Antigenicity Against BT21

Some of the identified epitopes arose due to unexpected cancer-specific gene regulatory events. For instance, we identified an epitope derived from TBC1D16, which was generated from the canonical *TBC1D16-201* transcript and a transcript arising from an alternative transcription start site, i.e., *TBC1D16-202*. When assessing recognition of other HLA-A*24 *TBC1D16*-expressing tumor cell lines (HepG2, PC3, and HT-144), only HT-144 was recognized by the TCR. Analysis of the sequence reads mapping to *TBC1D16* revealed that only BT21 and HT-144 contain the alternative *TBC1D16-202* transcript (**Figure 4A**). Overexpression of the canonical isoform of *TBC1D16-201* and the transcriptional variant *TBC1D16-202* in HLA-A*24:02 293T cells led, in both cases, to T cell activation. However, based on the average number of transgene integrations per APC and the elicited GFP+ activation signal, a much stronger GFP+ signal was observed for TBC1D16-202 than TBC1D16-201, suggesting that the TBC1D16-202-derived epitope is predominantly presented (**Figure 4B**). Remarkably, the transcriptional variant *TBC1D16-202*, which leads to a shorter TBC1D16 protein (47kDa), is induced by demethylation during melanoma tumorigenesis and promotes melanoma growth both *in vitro* and *in vivo*^31, 32^. Interestingly, while total *TBC1D16* expression was readily detectable in normal tissues (>10 TPM) and only modestly elevated in ∼50% of skin and uveal melanomas, quantification of the *TBC1D16-202* transcript provided markedly greater tumor discrimination, with >80% of melanomas expressing this isoform above the highest 95th-percentile expression observed in normal tissue (**Figure 4C**).

**Figure 4.**
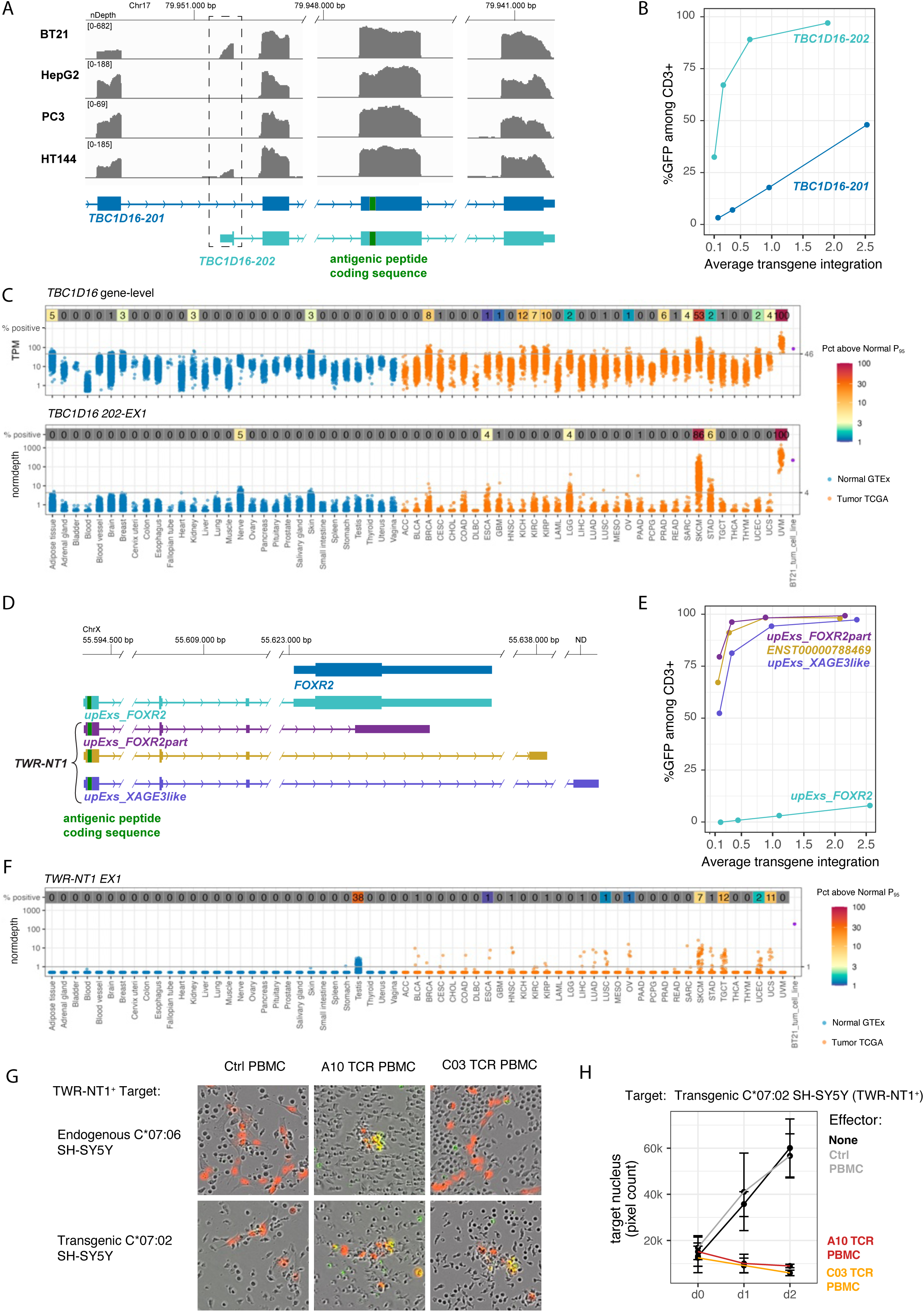
Complex Regulatory Events Underlying Tumor Antigenicity Against BT21. (A) Expression of *TBC1D16* transcript variants in 4 tumor cell lines (BT21, HepG2, PC3, HT144). Coverage plots reflect normalized sequencing depth scaled to 10 million mapped bases per sample. The Y-axis range reflects sequencing depth for each track. The lower panel shows the gene models for the 2 annotated transcript variants, *TBC1D16-201* and *TBC1D16-202*. The dashed box highlights the alternative transcription start region specific to the *TBC1D16-202* isoform. The genomic region encoding the presented epitope is indicated in green. Genomic coordinates are shown above the coverage tracks. (B) Relationship between *TBC1D1*6 isoform overexpression and T cell activation. HLA-A*24:02 293T transduced with the coding sequence (CDS) of *TBC1D16-201* and *TBC1D16-202* isoforms linked to a RFP construct at different dilutions were co-cultured with reporter T cells transduced with the TCR recognizing the *TBC1D16*-derived epitope (B04). RFP positivity was used to determine transgene integration. Cells were stained with anti-CD3 antibodies, and the percentage of GFP+ cells among CD3+ T cells was measured for each condition. (C) Expression of *TBC1D16* transcripts across normal tissues from the Genotype-Tissue Expression (GTEx, blue) portal and tumors from The Cancer Genome Atlas (TCGA, red) are shown. The upper panel depicts gene-level expression in transcripts per million (TPM, log scale), whereas the lower panel depicts genomic locus-based expression quantified from normalized RNAseq coverage. Each dot represents an individual sample. Numbers above tissue and cancer types indicate the percentage of samples with expression above the 95th percentile of normal tissues (P_95_), testis excluded. Cancer types are abbreviated according to TCGA nomenclature. SKCM and UVM stand for skin and uveal melanoma. (D) Schematic representation of the genomic locus containing *TWR-NT1* related transcripts. Exons represent boxes, introns are labeled as arrows, with taller boxes indicating the CDS. The genomic region encoding the antigenic peptide is highlighted in green. Scale bars show genomic coordinates. (E) Relationship between overexpression of *TWR-NT1* transcripts and T cell activation. HLA-C*07:02 293T were transduced with CDS of 4 *TWR-NT1* related transcripts coupled to RFP at different dilutions and were co-cultured with reporter T cells transduced with the TCR of interest (A10). RFP positivity was used to determine transgene integration and plotted against the percentage of GFP+ cells among CD3+ T cells. (F) *TWR-NT1* expression across normal tissues from GTEx (blue) and tumors from TCGA (orange). For each gene, the Y-axis shows expression in TPM (log scale), and each dot represents an individual sample. The numbers and color above each tissue or cancer type indicate the percentage of samples with expression above the 95th percentile of normal tissues (P_95_), excluding testis. Cancer types are abbreviated according to TCGA nomenclature. SKCM and UVM stand for skin and uveal melanoma. (G,H) Functional validation of the TWR-NT1 derived epitope in the neuroblastoma cell line SH-SY5Y. SH-SY5Y cells transduced with HLA-C*07:06 or HLA-C*07:02 and expressing RFP are co-cultured with healthy donor PBMCs expressing TCRs targeting the TWR-NT1 epitope (A10 and C03) or with untransduced control PBMCs. (G) Representative Incucyte images show tumor cell growth (red nucleus) and T cell-mediated cytotoxicity (caspase 3/8 green dye). (H) Longitudinal quantification of tumor burden based on target nucleus counts. Cultures containing TCR-transduced PBMCs exhibited sustained suppression of tumor growth compared with control conditions. Measurements were performed in duplicate. Error bars represent the 95% confidence interval.

Similarly, the TWR-NT1 epitope is derived from a group of transcripts that overlap with *ENSG00000302645*, *FOXR2* and a XAGE3-like locus on chromosome X (**Figure 4D**). *FOXR2* is annotated as a single-exon gene, but multiple upstream exons have been reported in tumors^33, 34^. Specifically, this epitope is recognized by two distinct TCRs and originates from a 132-bp ORF spanning two of these upstream exons. However, transgenic expression of a transcript composed of these exons in C*07:02 293T cells resulted in only weak T cell activation (**Figure 4E**). When analyzing BT21-derived RNA by full-length direct RNA sequencing, we identified and cloned 3 transcripts containing the antigenic sequence, which were all composed of upstream exons spliced with a downstream exon consisting of either a portion of *FOXR2* exon, a distal exon (∼15kb away) corresponding to *ENST00000788469*, or an unlocalized *XAGE3*-like locus. All these transcripts individually elicited strong T cell activation, suggesting that the ORF encoding the epitope is primarily translated when it is not in competition with the full *FOXR2* CDS (**Figure 4D,E**). Notably, the upstream exons exhibited cancer testis-like expression, being undetectable in normal tissues except testis, while overexpressed in 7% of skin melanomas and 11% of uterine carcinosarcomas (**Figure 4F**). When incubating the neuroblastoma cell line SH-SY5Y, which also expresses the upstream exons, with primary T cells transduced with TWR-NT1 specific TCRs, we observed tumor cell killing and sustained growth suppression. C03 TCR was strictly dependent on HLA-C*07:02, whereas the A10 TCR also recognized its peptide presented on the SH-SY5Y endogenously expressed HLA-C*07:06 (**Figure 4G,H**), suggesting that certain TCRs can cross-recognize their epitope across different HLA alleles.

Finally, we also identified an epitope encoded by a canonical transcript and an alternative *ALKBH7-202* transcript that results from an alternative transcriptional start site. Both transcripts are expressed in BT21, but the *ALKBH7-202* transcript yields more pronounced T cell activation compared to the canonical transcript (**Figure S4A,B**). Interestingly, the canonical ALKBH7 protein is targeted to the mitochondrial matrix via an N-terminal targeting signal that is absent in the shorter *ALKBH7-202* transcript. This different subcellular localization of the transcript most likely explains why *ALKBH7-202* triggers stronger T cell activation.

### TWISTAR Deorphanizes TCRs Clonally Expanding during Checkpoint Immunotherapy

Next, we assessed TWISTAR’s ability to deorphanize TCRs that clonally expand in patients receiving ICB. We selected a patient from a cohort of treatment-naïve early breast carcinoma patients receiving a single dose of anti-PD1 before undergoing tumor resection^1^. Based on single-cell profiling of paired pre- and on-treatment biopsies, the 12 most abundant CD8+ clonotypes in the on-treatment biospy were selected: 10 TCRs showed a ≥3-fold increase in relative frequency between on-*versus* pre-treatment biopsies, indicating that these clonotypes expanded during ICB (**Figure 5A**). Most of the T cells carrying these clonotypes were annotated as exhausted T cells, exhibiting an activated phenotype (*PRF1*, *GZMB*) with exhaustion-like features (*PD1*, *LAG3*) and prominent *CXCL13* expression^26^ (**Figure S5A,B**). T cells containing the other 2 non-expanding TCRs (i.e., T02 and T07) clustered as naïve/central-memory or resident-memory T cells and did not increase in relative frequency. TWISTAR identified cognate antigens for 7 out of 10 expanded clonotypes, but not for the non-expanded clonotypes (**Figure 5B,C**). The 7 expanding TCRs recognized 5 tumor antigens, for which the predicted epitopes were confirmed in peptide pulse experiments (**Figure S5C,D**). None were derived from somatic mutations: one originated from a canonical CDS in *FTHL17* (2 TCRs), while 3 arose from a novel CDS within known transcripts, i.e., *RAB11B* (2 TCRs), *EXOSC3* and *PRPF6*. In case of *RAB11B* and *EXOSC3*, both epitopes were derived from upstream ORFs. Using the Immunogenic Search Engine^10^, which identifies both canonical and non-canonical antigens from public immunopeptidomics datasets, we detected peptides derived from the same ORF in *RAB11B* and *EXOSC3* (i.e., in an acute myeloid leukemia and 3 melanoma samples, respectively; **Supplementary Table 1**). Finally, one epitope was derived from an intronic sequence in *HELZ*. Inspection of sequencing reads derived from the tumor’s transcriptome revealed that read density across the intron was irregular, suggesting existence of an unannotated *de novo* transcribed region within this intron (**Figure S6A**).

**Figure 5.**
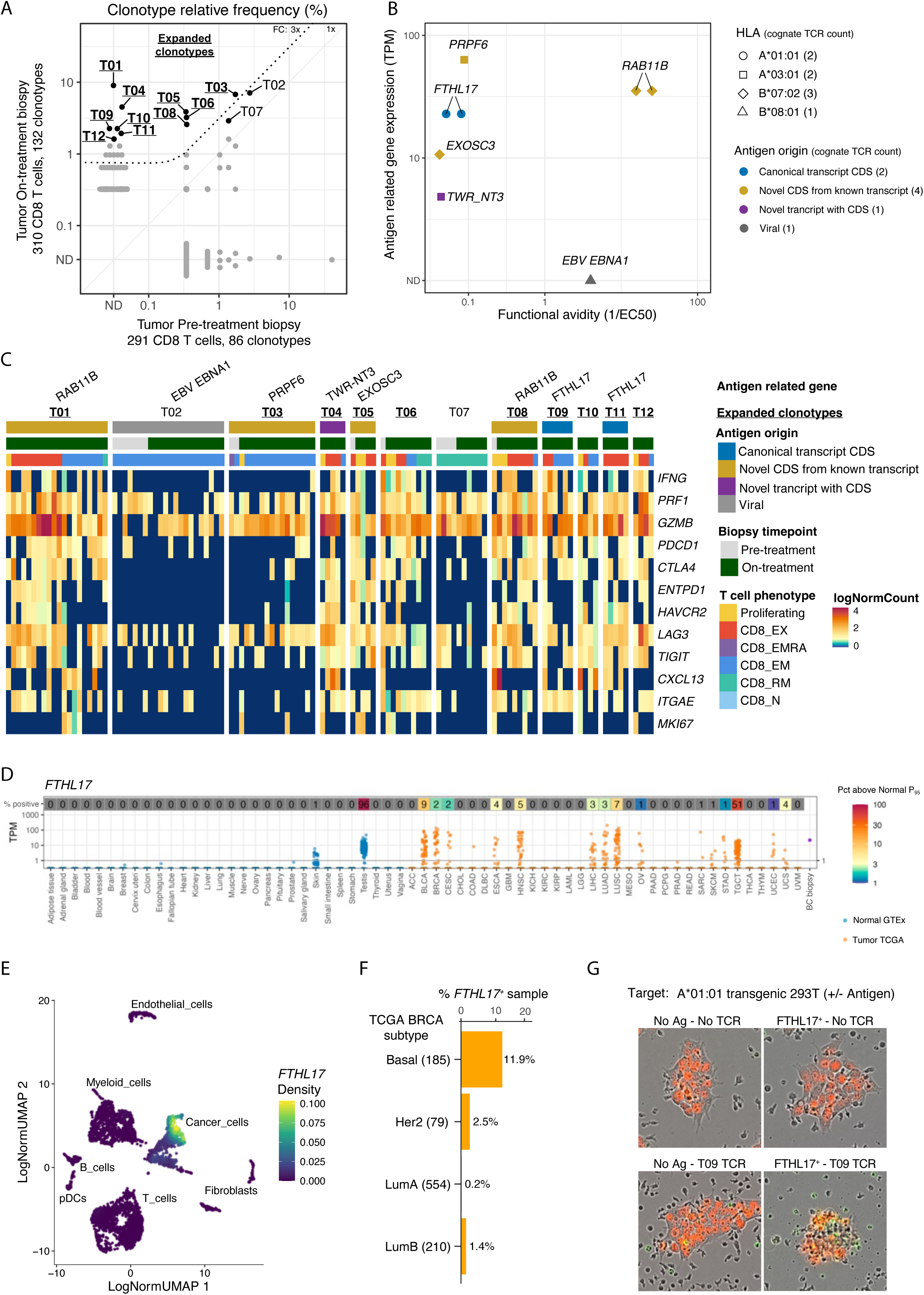
Deorphanization of TCRs Clonally Expanding during Checkpoint Immunotherapy. (A) Clonotype relative frequency of selected T cells in pre- and on-treatment tumor biopsies from a breast cancer patient. The relative frequency of 86 and 132 CD8+ T-cell clonotypes is shown. Expanded clonotypes are highlighted in bold and underlined. A dotted line indicates a 3-fold expansion threshold and at least 3 cells detected in the on-treatment biopsy. (B) Characteristics of antigens recognized by deorphanized TCRs from breast cancer tumor-infiltrating lymphocytes (TILs). Each dot represents the cognate antigen of one deorphanized TCR. The graph depicts functional avidity defined as the inverse of the half-maximal effective concentration (EC_50_, µM) relative to antigen-related gene expression determined by bulk RNAseq data. HLA restriction of each TCR–antigen pair is indicated by a symbol, while the type of transcript from which the antigen is derived is indicated in color. (C) Single-cell gene expression heatmap with rows representing individual genes and columns representing individual T cells, grouped based on antigen specificity, clonotype frequency, biopsy timepoint and T cell phenotype. Cognate antigens are indicated on the top. (D) *FTHL17* expression across normal tissues from the Genotype-Tissue Expression (GTEx, blue) portal and tumors from The Cancer Genome Atlas (TCGA, orange) are shown. For each gene, the Y-axis shows expression in transcripts per million (TPM, log scale), and each dot represents an individual sample. The numbers and color above each tissue or cancer type indicate the percentage of samples with expression above the 95th percentile of normal tissues (P_95_), excluding testis. Cancer types are abbreviated according to TCGA nomenclature. BRCA, BLCA, LUSC and HNSC represent breast, bladder, small cells lung, and head and neck cancer, respectively. (E) UMAP visualizing *FTHL17* expression in single-cell data of pre- and on-treatment biopsies from a breast cancer patient. Cells are colored according to normalized *FTHL17* expression density. Major cell populations are indicated on the UMAP, including cancer cells, T cells, B cells, myeloid cells, endothelial cells, pDCs, and fibroblasts. (F) Prevalence of *FTHL17* expression in TCGA breast cancer molecular subtypes. Bars indicate the percentage of samples with an *FTHL17* expression level above 1 TPM. Samples were considered positive when *FTHL17* expression exceeds a predefined expression threshold (see **Methods**). Numbers in parentheses denote the number of tumors per subtype, while subgroup percentages are shown next to each bar. (G) Functional validation of the FTHL17-derived epitope in HLA-A*01:01-expressing 293T cells stably expressing the *FTHL17* full-coding sequence (or no sequence, as control) and a downstream RFP cassette. Cells were co-cultured with healthy donor PBMCs transduced with the TCR recognizing the FTLH17-derived epitope (T09) or with untransduced control PBMCs. Representative Incucyte images show growth of target cells (red nucleus) and T cell-mediated cytotoxicity (AnnexinV green dye).

Tumor-infiltrating T cells that are non-reactive towards tumor antigens are referred to as bystanders and often recognize viral-derived epitopes^35^. Therefore, we screened the 5 TCRs that did not identify a tumor antigen against a library of 108,603 sequences tiling the human virome^15^. This identified a single non-expanding TCR specific for an Epstein-Barr virus (EBV)-derived epitope presented on HLA-B*08:01 (**Figure 5B,C**). Notably, The Immune Epitope Database (IEDB) reports the exact same TCR (receptor group ID 4934) to be specific for a distinct cytomegalovirus (CMV)-derived peptide presented on HLA-A*03:01 based on pMHC tetramer data. However, we observed no reactivity towards this CMV peptide when pulsed onto cells expressing HLA-A*03:01 or HLA-B*08:01 (**Figure S5D**), suggesting that the reported association of this TCR with the CMV peptide is erroneous. These findings highlight a limitation of database-driven TCR-antigen assignment and demonstrate that TWISTAR can also reliably identify virus-associated antigens.

Interestingly, although *FTHL17* was described as a cancer testis gene >20 years ago^36^, our findings provide the first evidence of T cell recognition of a FTHL17-derived epitope. We identified two distinct TCRs recognizing the same HLA-A*01:01-restricted epitope derived from FTHL17. Both TCRs were undetectable in the pre-treatment biopsy but expanded to represent >4% of CD8+ T cells in the on-treatment sample (0/291 *versus* 6/310 and 7/310 CD8⁺ T cells, respectively). Notably, although both clonotypes express markedly different TCR β-chains, their α-chains were highly similar (**Supplementary Table 2**). Both TCRs also had comparable functional avidities, suggesting that their antigen binding predominantly relies on the α chain. In contrast to 3 of the 4 other antigen-coding genes (**Figure S6B**), *FTHL17* transcript expression was undetectable in normal tissue (besides testis and a few skin samples), but upregulated in various cancer types, including bladder, lung, and head and neck carcinomas (**Figure 5D**). Furthermore, single-cell transcriptomic analysis demonstrated that *FTHL17* expression was specific to cancer cells within the patient’s tumor (**Figure 5E)**. Notably, within breast cancer datasets, *FTHL17* expression was enriched in the more aggressive basal subtype, where it was detected in 11.9% of cases (**Figure 5F**). Co-culturing HLA-A*01:01 293T cells transfected with an FTHL17-encoding plasmid with primary T cells transduced with an FTHL17-specific TCR confirmed potent cytolytic activity (**Figure 5G, S6C,D**).

### Immune Dominance Underlying a Complete Response to ICB

Finally, we analyzed the immune infiltrate of an advanced HCC patient, who initially presented with a 12-cm tumor lesion. Following 2 months of ICB therapy, the patient achieved a profound radiological response, resulting in post-treatment resection of the residual lesion. Histopathological evaluation of the resection specimen demonstrated a complete pathological response, with no viable tumor cells remaining. This exceptional response was associated with long-term survival, providing a unique opportunity to investigate the tumor antigens driving this effective anti-tumor immune response. Both pre- and post-treatment resection specimen were analyzed at single-cell level. Despite the 2-month interval between biopsy and resection, we found that most of the dominant clonotypes present prior to treatment persisted in the residual lesion. We selected the 12 most frequent CD8+ T cell clonotypes from both lesions; 5 clonotypes were shared between both samples, resulting in a total of 19 TCRs (**Figure 6A, S7A,B**).

**Figure 6.**
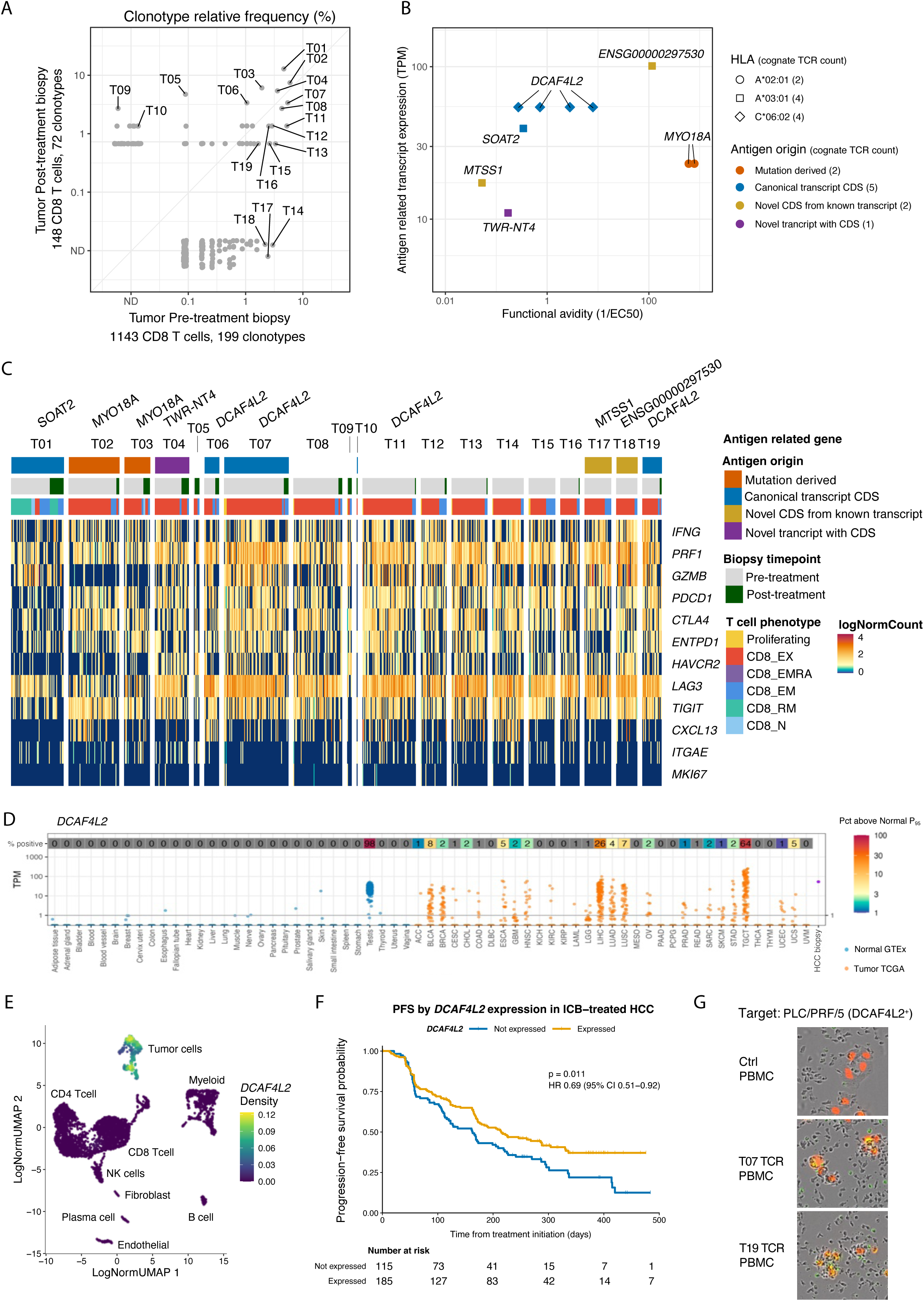
Immune Dominance Underlying a Complete Response to ICB in an HCC patient. (A) Clonotype relative frequency of selected T cells in pre- and post-treatment biopsies from a hepatocellular carcinoma (HCC) patient. The relative frequency of 199 and 72 CD8+ T-cell clonotypes is shown. Expanded clonotypes are highlighted in bold and underlined. A dotted line indicates a 3-fold expansion threshold and at least 3 cells detected in the post-treatment biopsy. (B) Characteristics of antigens recognized by deorphanized TCRs from tumor-infiltrating lymphocytes (TILs) in an HCC patient. Each dot represents the cognate antigen of one deorphanized TCR. The graph depicts functional avidity defined as the inverse of the half-maximal effective concentration (EC_50_, µM) relative to antigen-related gene expression determined by bulk RNAseq data. HLA restriction of each TCR–antigen pair is indicated by a symbol, while the type of transcript from which the antigen is derived is indicated in color. (C) Single-cell gene expression profile of TILs with rows representing individual genes and columns representing individual T cells, grouped based on antigen specificity, clonotype frequency, biopsy timepoint and T cell phenotype. Cognate antigens are indicated on the top. (D) *DCAF4L2* expression across normal tissues from the Genotype-Tissue Expression (GTEx, blue) portal and tumors from The Cancer Genome Atlas (TCGA, orange). For each gene, the Y-axis shows expression in transcripts per million (TPM, log scale), and each dot represents an individual sample. The numbers and color-coded annotations above each tissue or cancer type indicate the percentage of samples with expression exceeding the 95th percentile of normal tissue expression (P_95_), excluding testis. Cancer types are abbreviated according to TCGA nomenclature. (E) UMAP visualizing *DCAF4L2* expression in single-cell data of pre- and post-treatment biopsies from an HCC patient. Cells are colored according to normalized *DCAF4L2* expression density. Major cell populations are indicated on the UMAP, including cancer cells, T cells, B cells, myeloid cells, endothelial cells, pDCs, and fibroblasts. (F) Kaplan-Meier plot of progression-free survival in 300 patients with HCC treated with atezolizumab-based therapy stratified according to baseline tumor *DCAF4L2* expression. The hazard ratio was estimated using a Cox proportional hazards model, and the P value using the log-rank test. (G) Functional validation of the DCAF4L2-derived epitope in the *DCAF4L2*+ HCC cell line PLC/PRF/5. Target cells were modified to express HLA-C*06:02 and RFP and co-cultured with healthy donor PBMCs transduced with TCRs (T07 or T19) recognizing the DCAF4L2-derived epitope or untransduced control PBMCs. Representative Incucyte images show growth of target cells (red nucleus) and T cell-mediated cytotoxicity (AnnexinV green dye).

Overall, TWISTAR deorphanized 10 TCRs, corresponding to 6 distinct tumor antigens (**Figure 6B, S7C,D**): one was derived from a somatic mutation in *MYO18A*, 2 from a canonical CDS in *DCAF4L2* and *SOAT2*, 2 from a novel CDS in previously described transcripts, i.e., an alternative ORF in *MTSS1* and an ORF in the lncRNA encoding transcript *ENSG00000297530*, and finally, one was derived from a locus annotated as a *VNN2* intron. Peptide pulse experiments revealed that the epitopes derived from the *MYO18A* mutation and lncRNA *ENSG00000297530* exhibited the highest EC_50_ values. Unexpectedly, in case of *MYO18*, the corresponding wild-type peptide was also capable of activating the cognate TCR, although with reduced potency, suggesting that the mutation enhances peptide generation and/or HLA presentation efficiency rather than creating a new TCR recognition motif (**Figure S7D**). In TCGA, expression of the lncRNA *ENSG00000297530* was observed predominantly in lung, liver and colorectal carcinomas (7%, 5%, and 3% of cases, respectively), while normal tissue expression was largely absent except for low-level signals in lung and testis (**Figure S8B**). In the single-cell data of the patient’s tumor, *ENSG0000029753* expression was specifically observed in the cancer cell subcluster (**Figure S8C**). The most frequent T cell clonotype in the resection specimen (12.8% of CD8 T cells) recognized an epitope derived from *SOAT2*. In TCGA, 11% of HCC tumors exhibited *SOAT2* expression at higher levels than in normal tissue (**Figure S8A**). In the single-cell data *SOAT2* expression was also confined to cancer cells (**Figure S8C**).

Finally, we also identified 4 TCRs recognizing 2 overlapping DCAF4L2-derived epitopes (NYCRIAREL and CRIARELRV), both presented on HLA-C*06:02. Three of the T cell clonotypes were detected in both biopsies at frequencies ranging between 1-10%, together representing 8.1% and 8.8% of the TCRs detected in the pre- and post-treatment biopsy, respectively (**Figure 6C**). A fourth clonotype emerged only in the resection specimen. Remarkably, CDR3 sequences of α- and β-chains of TCRs recognizing both DCAF4L2-derived epitopes shared minimal homology, indicating diverse binding modalities. To assess whether DCAF4L2-derived epitopes could also be presented on other HLA alleles and thus broaden the patient population that might benefit from targeting this antigen, we generated a TCR library comprising all TCR sequences identified in the paired biopsy samples and screened it against DCAF4L2-expressing 293T cells presenting either HLA-C*06:02 or HLA-A*02:01. The latter was selected because it represents one of the most prevalent HLA-A alleles worldwide. On HLA-C*06:02, the four previously identified DCAF4L2-specific TCRs were recovered, together with two additional TCRs recognizing the same two overlapping epitopes. On HLA-A*02:01, two additional DCAF4L2-specific TCRs were identified, both recognizing a different epitope derived from a distal region of the DCAF4L2 coding sequence (**Figure S8D,E**). Thus, a single shared tumor-associated antigen can engage diverse TCR clonotypes through multiple epitopes presented by distinct HLA alleles. This highlights that such antigens may support broader immune responses than individual mutation-derived neoantigens, which are often restricted to a single mutant peptide-HLA combination. *DCAF4L2* exhibits testis-restricted expression in normal tissue and has been implicated in promoting invasion and metastasis in colorectal cancer through functional perturbation studies^37^. In TCGA, *DCAF4L2* is detected in several cancer types, including bladder, lung and esophageal carcinoma, but is most frequently expressed in HCC (26%; **Figure 6D**). In the patient’s single cell data, *DCAF4L2* expression was confined to the cancer cell compartment (**Figure 6E**). Overall, these data strongly suggest that DCAF4L2-derived epitopes act as major drivers of the patient’s immune response against this tumor.

### Shared DCAF4L2 antigens drive ICB-induced immune responses in HCC

To further explore the role of DCAF4L2 in mediating tumor regression during ICB, we analyzed bulk RNAseq data from the IMbrave150 trial, which evaluated the addition of atezolizumab (anti-PD-L1) to bevacizumab (anti-VEGF) *versus* sorafenib (multi-kinase inhibitor) for the treatment of patients with unresectable HCC^38^. Stratification based on *DCAF4L2* expression revealed that *DCAF4L2+* patients treated with ICB exhibited improved progression-free survival relative to patients lacking *DCAF4L2* expression (HR=0.69; 95%CI 0.51-0.92, **Figure 6F**), while such effects were not observed in patients treated with sorafenib as a control (**Figure S8F**). This suggests that DCAF4L2 may be a common driver of response to ICB in advanced HCC. To confirm that DCAF4L2 results in functional antigen presentation and recognition of tumor cells by T cells, we expressed two DCAF4L2-specific TCRs in primary T cells and co-cultured these with the *DCAF4L2+* PLC/PRF/5 hepatoma cell line. This resulted in targeted cytolysis and sustained growth suppression (**Figure 6G, S8G**), thereby confirming DCAF4L2 as an actionable tumor antigen that enables potent T cell-mediated killing.

## DISCUSSION

Here, we developed TWISTAR, a novel screening platform that enables the systematic deorphanization of selected TCRs against a library of autologous tumor-derived cDNA. A unique aspect of TWISTAR is that it uses single-cell expression and TCR data as input to select relevant TCR clonotypes and then identifies their cognate antigens. By selecting TCRs from T cells that are tumor-reactive against an autologous tumor cell line, or that clonally expand under ICB and express both exhausted and activated markers, TWISTAR reliably identifies tumor antigens driving anti-tumor responses. This approach thus provides an unbiased and scalable framework for systematically mapping the functional antigen landscape directly from patient tumors.

A key finding is that neoantigens account for only a minority of the antigens recognized by tumor-infiltrating T cells in our datasets (i.e., 2 neoantigens identified in BT21, 1 in HCC, and none in breast cancer). Instead, the dominant fraction of epitopes originated from non-mutated sequences, including canonical tumor-associated antigens, as well as many non-canonical transcriptional and translational events. This observation suggests that the immunogenic landscape of tumors extends far beyond antigens predicted to arise from somatic mutations in coding sequences. This is relevant as neoantigen-focused tumor vaccination approaches hold great therapeutic promise but suffer from limitations. Particularly, most computationally predicted neoantigens included in vaccines turn out to be non-immunogenic, leading to a considerable fraction of patients having either no immune response, or responding to only a limited subset of vaccine-encoded epitopes^39^. Since tumor vaccination appears to be clinically most impactful when at least 3-4 antigens trigger an anti-tumor response^40^, there is a critical need to identify additional tumor antigens that are truly immunogenic. TWISTAR addresses these issues identifying antigens derived from non-mutated transcripts, and, inherent to its underlying methodology, these antigens are recognized by active T cells that have triggered an immune response in the patient. While efforts with immunopeptidomics have already highlighted that many MHC-I presented peptides arise from non-canonical events^41^, our study provides evidence that a subset of these antigens is effectively recognized by T cells and drives tumor-reactive immune responses in patients. The observation that several of the underlying non-canonical transcripts have also been linked to oncogenic or metastasis-promoting functions further suggests that these antigens do not represent stochastic noise, but rather stable targets expressed in a high fraction of cancer cells; properties that render them therapeutically exploitable.

Intriguingly, many of the detected antigens are derived from transcripts that are shared across patients and cancer types, but are absent or weakly expressed in normal tissues, except for immune-privileged sites such as the testis, a profile reminiscent of classical cancer testis antigens. This expression pattern has been associated with DNA hypomethylation and epigenetic reactivation of germline programs in cancer^42, 43^. However, unlike classical cancer testis antigens such as NY-ESO-1, FTHL17 and DCAF4L2 appear to be expressed across multiple cancer types but only rarely in melanoma. This suggests that these antigens may not conform to the classical melanoma-associated cancer testis antigens expression profile and could instead be subject to distinct regulatory mechanisms. Nevertheless, the high recurrence of these transcripts across malignancies points to the existence of a shared tumor antigenome, in which diverse cancers converge on a relatively limited set of immunogenic transcriptional programs. This has important implications for immunotherapy, as such shared antigens may enable the development of broadly applicable T cell-based interventions, including off-the-shelf vaccines, which offer a highly advantageous alternative to personalized neoantigen vaccines in terms of cost and scalability. Their effectiveness, however, would depend on HLA compatibility between the identified antigens and the patient’s HLA type. Encouragingly, since most transcripts identified by TWISTAR contain ORFs of several hundred nucleotides that are specifically expressed in tumors, these ORFs are likely to generate additional epitopes presented by other HLA alleles. Indeed, we already identified NY-ESO-1 and DCAF4L2-derived epitopes presented by distinct HLA alleles and recognized by different TCRs. Importantly, within this shared antigen landscape, we also observed evidence of immune dominance, whereby multiple clonally expanded T cell populations converge on a limited number of antigenic peptides derived from the same transcripts. In breast cancer, we identified 2 distinct TCRs directed against the same peptide from FTLH17 emerging during ICB, whereas in HCC there were 8 divergent TCRs directed against 3 distinct peptides from DCAF4L2 presented by two different HLAs. Such convergence suggests that anti-tumor immune responses are not evenly distributed across all available antigens but are instead shaped by a restricted hierarchy of immune dominant epitopes that guide the effectiveness of ICB.

Interestingly, while neoantigens are often assumed to generate high-affinity TCR responses due to their non-self nature and consequent lack of thymic deletion, tumor-associated antigens being derived from self-proteins, are thought to elicit weaker responses due to negative selection in the thymus. However, our peptide pulse experiments revealed that functional TCR avidity does not strictly follow this expected hierarchy based on antigen class. We observed substantial heterogeneity in avidity comparing neoantigens with the other antigens identified. For instance, several non-canonical antigens derived from lncRNAs elicited unexpectedly high functional avidities, in some cases exceeding those observed for neoepitopes. These observations suggest that antigen processing efficiency, translational context, and subcellular protein localization may be as important as central tolerance in shaping the functional fitness of tumor-reactive T cell repertoires.

As an antigen discovery platform, TWISTAR provides a broadly accessible method that complements existing antigen discovery strategies, without the need for specialized equipment beyond standard cell culture facilities and automated microscopy. Unlike computational neoantigen prediction, which is limited by prediction accuracy and incomplete modeling of antigen processing, or immunopeptidomics, which identifies presented peptides without establishing immunogenicity, TWISTAR directly measures T cell recognition. Compared to proteome-wide or random peptide display systems, such as TSCAN^15^ and TCR-MAP^16^, TWISTAR preserves biological relevance by restricting the antigen search space to antigens naturally processed in patient tumors. On the other hand, since it screens against a cDNA-derived library of tumor transcript, it also detects unannotated and non-canonical transcripts, which significantly expands the breadth of antigens discovered. Compared to these other platforms, which so far deorphanized only a handful of TCRs, TWISTAR is a highly scalable platform, as evidenced by the deorphanization of up to 40 TCRs. Despite these advances, certain limitations remain. TWISTAR relies on *in vitro* co-culture systems that, while capturing key aspects of antigen processing and presentation, may not fully recapitulate the tumor microenvironment, including post-translational modifications, metabolic context, or spatial antigen sequestration within the tumor microenvironment. In addition, low tumor content or clonal heterogeneity may reduce the representation of cancer cell-derived transcripts during cDNA library preparation. Additionally, while TWISTAR effectively identifies antigens presented by HLA class I molecules, the extension to CD4⁺ T cells and HLA class II-restricted epitopes remain unexplored.

Beyond these limitations, our findings position TWISTAR as a highly effective approach for the systematic exploration of tumor antigen specificity. Applying TWISTAR to T cells derived from tumors with an activated immune compartment highlights a previously underappreciated breadth of immunogenic tumor antigens arising from diverse transcriptional programs. More broadly, our results support a unifying model in which tumor immunogenicity emerges from a composite antigenome consisting of mutations, canonical tumor-associated proteins, and a large and previously underexplored layer of non-canonical transcriptional products. The immune system appears to preferentially target a constrained subset of this landscape, defined by antigen processing efficiency, expression patterns, and HLA compatibility, rather than by mutational status alone. This framework challenges the prevailing neoantigen-centric view of tumor immune recognition and provides a roadmap for systematically incorporating non-canonical antigens into next-generation immunotherapies, from vaccines to TCR-engineered T cells, with the potential to reach a substantially broader patient population.

## RESOURCE AVAILABILITY

### Lead contact

Requests for further information and resources should be directed to and will be fulfilled by the lead contact, Diether Lambrechts.

### Materials availability

All unique/stable reagents generated in this study are available from the lead contact upon completion of a material transfer agreement, unless third party MTA restrictions apply that do not allow the direct sharing of materials.

### Data and code availability

Single-cell RNA-seq and TCR-seq data have been deposited at National Center for Biotechnology Information BioProject with accession code PRJNA985415 and at the European Genome-phenome Archive as EGAS00001004809 and EGAD50000001227. Requests for accessing raw sequencing data will be reviewed by the VIB data access committee. Any data shared will be released via a Data Transfer Agreement that will include the necessary conditions to guarantee protection of personal data according to European GDPR law. Read count data for the breast cancer and HCC patient are also available at https://lambrechtslab.sites.vib.be/en/dataaccess as of the date of publication. All other data reported in this paper will be shared by the lead contact upon reasonable request. Any additional information required to reanalyze the data reported in this paper is available from the lead contact upon request.

## ACKNOWLEDGMENTS

This work has been supported by an ERC advanced grant EXPAND-IT (grant no. 101055422), ERC Proof of concept TWISTAR (grant no 101335284)., Research Foundation Flanders FWO (grant no. G063727N, G093821N), Stichting Tegen Kanker (2024-161), KU Leuven internal fund (grant no. C14/18/092) to D.L. M.P is supported by an ERC grant CENTRIC-BRAIN (grant no. 101141901). J.D. is supported by a Senior Clinical Investigator mandate from the FWO (grant no. 1800926N). A.S. is supported by Fonds Nadine De Beauffort and by a Kom op Tegen Kanker grant. The computational resources and services used in this work were partly provided by the Flemish Supercomputer Centre (VSC), funded by the Research Foundation Flanders (FWO) and the Flemish Government department EWI. Funding to pay the Open Access publication charges for this article was provided by institutional funding (VIB).

## AUTHOR CONTRIBUTIONS

O.B., J.D. and D.L. conceived and designed the study. O.B. and D.L. wrote the manuscript based on all author’s contributions. O.B. and D.L. supervised the work. O.B., F.P., R.S.R., G.H., J.vd.R, E.W.G., J.D. and D.L. discussed and interpreted the data. O.B., R.S.R., J.X. and G.P. designed and performed bioinformatic analyses. E.L., A.S., M.P. and E.W.G. contributed data or collected human samples and related clinical data for the study. O.B., C.L., T.V.B., E.V. and R.S. performed TWISTAR and other related experiments or provided technical and experimental support.

## SUPPLEMENTAL FIGURE LEGENDS

Figure S1. Optimization and Validation of TWISTAR

(A, B) Fluorescence micrographs showing GFP+ cluster generation using 2D3 (A) and SL8 (B) reporter T cell lines, transduced with a low-affinity TCR recognizing the mutPIK3CA antigen expressed by all RFP+ co-cultured antigen-presenting cells (APCs).

(C) Flow cytometry-based analysis of GFP signal in 2D3 (control) and 48 clones of Jurkat-E6.1-derived NFAT-GFP reporter cells expressing a low-affinity mutPIK3CA-specific TCR and CD8 after co-culture with APCs expressing a mutPIK3CA-encoding minigene. Median fluorescence intensity (MFI) of GFP+ cells and frequency of GFP+ cells among CD3+ T cells are shown. The SL8 clone is highlighted in blue.

(D) Surface TCR expression measured by flow cytometry in reporter T cells transduced with the indicated retroviral or lentiviral vectors encoding the low-affinity mutPIK3CA-specific TCR.

(E) HLA-I surface expression measured by flow cytometry on 293T HLA-I knockout cells overexpressing HLA-A*03:01 and the indicated peptide-loading complex components.

(F) Effect of peptide-loading complex overexpression in APCs on the activation of SL8 reporter T cells expressing the low-affinity mutPIK3CA TCR. After overnight co-culture, CD3+ cells were analyzed for GFP expression by flow cytometry. Relative activation is normalized to untransduced APCs. Errors bars represent 95% confidence intervals on two technical replicates, * indicates *p*<0.05 by Student’s t test.

(G) Comparison of RNAseq and TWISTAR library coverage at gene and single-base resolution across coding sequences (CDS), annotated transcript loci outside of CDS (other_ANN), loci within 1 kb of annotations (Near_from_ANN), and loci >1 kb from annotations (Far_from_ANN). Only expression >1 transcript per million (TPM) or >1 base per billion (BPB) is shown. Pearson correlation between RNAseq and TWISTAR is indicated at the center of each graph, followed by the number of elements considered. Bottom-right percentages indicate the fraction of loci with >3 TPM/BPB in high-depth RNAseq also detected in the TWISTAR library at >1 TPM/BPB. The Pearson correlation coefficient is indicated in the center of each graph together with the number of gene or locus measurements included in the analysis.

Figure S2. Peptide-pulse Experiments on Predicted Epitopes in the BT21 cell line

(A,B) Validation and characterization of BT21-derived epitopes. Single HLA-I expressing 293T cells were pulsed with the indicated peptides at varying concentrations for 2 h (A) or at 50 μM (B), washed, and co-incubated with TCR-transduced reporter T cells for 24 h. Peptide titration assays were used to determine EC_50_ values (A), whereas peptide truncation analysis defined the minimal epitope length supporting maximal TCR recognition (B). T-cell activation was assessed by measuring GFP expression using flow cytometry. Experiments are group per HLA class (HLA-A*24:02, HLA-B*07:02, HLA-C*03:03 and HLA-C*07:02) and the tested TCR is indicated above each experiment (e.g., A09, B04 and C22 for HLA-A*24:02). Measurements were performed in duplicate. Error bars represent the 95% confidence interval.

Figure S3. Mutation-derived and Tumor-associated Antigens recognized by BT21 TILs

(A,B) T cell activation in response to *MAGEA* constructs. (A) Mapping of putative immunogenic epitopes derived from MAGEA6. The *MAGEA6* transcript is shown and the cDNA (embedded within the full-length transcript in grey) that was identified in antigen-presenting cells (APCs) by GFP+ T cells is highlighted in green. Coding sequences (CDS) that were not recognized are indicated in grey (Trim a and Trim b), whereas the minimal antigenic CDS (Trim c, 207 nts) is highlighted in green. The epitope derived from this minimal CDS segment is mapped onto MAGEA family proteins, including MAGEA3, MAGEA6 and MAGEA12. Conserved and variable amino acid positions within the aligned peptides are shown, with residues contributing to HLA-A*24:02 binding indicated as anchor residues (green dots). (B) HLA-A*24:02 293T cells were transfected with full-length CDS of *MAGEA3*, *MAGEA6* or *MAGEA12*, or with truncated fragments of *MAGEA6* (Trim a, Trim b, Trim c). The percentage of GFP+ cells among CD3+ T cells after a co-culture with TCR-transduced (A04) reporter T cells is shown. Control conditions represent untransfected APC cells. Measurements were performed in duplicate. Error bars represent 95% confidence interval.

(C) Gene expression profiles of the antigen encoding sequences from deorphanized tumor-infiltrating lymphocytes (TILs) in normal tissues from the Genotype-Tissue Expression (GTEx, blue) portal and tumors from The Cancer Genome Atlas (TCGA, orange). For each gene, the Y-axis shows expression in transcripts per million (TPM, log scale) and each dot represents an individual sample. The numbers and color above each tissue or cancer type indicate the percentage of samples with expression above the 95th percentile of normal tissues (P_95_), excluding testis. Cancer types are abbreviated according to TCGA nomenclature.

(D) For transcripts not adequately represented by GENCODE v26 gene annotations, expression was quantified at the genomic locus level using normalized read depth derived from RNAseq coverage data. The same tissue and tumor comparison framework as in (C) is shown.

Figure S4. Characterization of the *ALKBH7* antigen-related locus

(A) Schematic representation of the genomic locus producing both *ALKBH7*-related transcripts. Transcripts are shown with exons as boxes and introns as arrows, with taller boxes indicating coding sequences (CDS). The genomic region encoding the antigenic peptide is highlighted in green. Scale bars indicate hg38 genomic coordinates.

(B) Relationship between *ALKBH7*-related transcript overexpression and T cell activation. HLA-A*24:02 293T cells were transduced with lentiviral constructs encoding ALKBH7-201 and ALKBH7-202 transcripts with a downstream RFP construct at different dilutions. RFP was quantified to determine transgene integration. After a co-culture with C22 TCR-transduced reporter T cells, the percentage of GFP+ cells among CD3+ T cells was compared with the transgene integration. Each point represents a distinct transduction level.

Figure S5. Single-cell data of the breast cancer patient selected for TCR deorphanization

(A) UMAP projection of tumor-infiltrating CD8+ T cells analyzed by single-cell profiling of tumor biopsies collected from a breast cancer patient. Cells on the UMAP are colored according to their phenotypic states: naïve (CD8_N), resident-memory (CD8_RM), effector-memory (CD8_EM), CD45_RA effector memory (CD8_EMRA), exhausted (CD8_EX) and proliferating T cells. Expanded clonotypes (T01–T12), which were deorphanized by TWISTAR, are shown in individual panels and projected onto the same UMAP.

(B) Expression of selected genes embedded on the UMAP. Feature plots show normalized expression of genes associated with effector function (*IFNG*, *PRF1*, and *GZMB*), inhibitory receptor or exhaustion programs (*PDCD1*, *CTLA4*, *ENTPD1*, *HAVCR2*, *LAG3* and *TIGIT*), tissue residency (*ITGAE*), dysfunctional or exhausted differentiation (*CXCL13*) and proliferation (*MKI67*). Colors indicate normalized gene expression levels (GEX).

(C,D) Validation and characterization of breast cancer-derived epitopes. Single HLA-I expressing 293T cells were pulsed with the indicated peptides at varying concentrations for 2 h (A) or at 50 μM (B), washed, and co-incubated with TCR-transduced reporter T cells for 24 h. Peptide titration assays were used to determine EC_50_ values (A), whereas peptide truncation analysis defined the minimal epitope length supporting maximal TCR recognition (B). Experiments are group per HLA class (HLA-A*01:01, HLA-A*03:01, HLA-B*07:02 and HLA-B*08:01) and the tested TCR is indicated above each experiment (e.g., T09 and T11 for HLA-A*01:01). T cell activation was assessed by measuring GFP expression using flow cytometry. Measurements were performed in duplicate. Error bars represent the 95% confidence interval.

Figure S6. Characterization of tumor antigens identified in the breast cancer patient

(A) RNAseq read coverage detected in the locus containing the *TWR-NT3* transcript on chromosome 17. Read depth is displayed in the upper track and individual sequencing reads are shown below. Gene annotations are shown at the bottom, including *HELZ* and *TWR-NT*3. The epitopes within TWR-NT3 are indicated in green. Genomic coordinates in hg38 are shown at the top.

(B,C) Expression of antigen-encoding genes and transcripts identified across normal tissues from the Genotype-Tissue Expression (GTEx, blue) portal and tumors from The Cancer Genome Atlas (TCGA, orange). For antigens represented by annotated genes, expression is shown in transcripts per million (TPM; log scale) (B). For antigens encoded by transcripts not represented in GENCODE v26, expression was quantified at the genomic locus level using normalized RNAseq read coverage (C). Each dot represents an individual sample. Numbers and colors above each tissue or cancer type indicate the percentage of samples with expression exceeding the 95th percentile of normal tissue expression (P_95_), excluding testis. Cancer types abbreviated according to TCGA nomenclature, BRCA represents breast cancer.

(D,E) Functional validation of the FTHL17-derived epitope in HLA-A*01:01-expressing 293T cells stably expressing the full *FTHL17* full coding sequence (or no sequence, as control) and a downstream RFP cassette. Cells were co-cultured with healthy donor PBMCs transduced with the TCR recognizing the FTLH17-derived epitope (T09) or with untransduced control PBMCs. (D) Representative fluorescent acquisition images corresponding to Figure 5G are shown for the indicated conditions, with growth of target cells (red nucleus) and T cell-mediated cytotoxicity (AnnexinV green dye). (E) Longitudinal quantification of images shown in panel D. Mean values are shown, error bars represent the 95% confidence interval.

Figure S7. Single-cell data of the hepatocellular carcinoma patient selected for TCR deorphanization

(A) UMAP projection of tumor-infiltrating CD8+ T cells analyzed by single-cell profiling. Cells in the UMAP are colored according to their phenotypic states: naïve (CD8_N), resident-memory (CD8_RM), effector-memory (CD8_EM), CD45_RA effector memory (CD8_EMRA), exhausted (CD8_EX) and proliferating T cells. Expanded clonotypes (T01–T12), which were deorphanized by TWISTAR, are shown in individual panels and projected onto the same UMAP.

(B) Expression of selected genes embedded on the UMAP. Feature plots show normalized expression of genes associated with effector function (*IFNG*, *PRF1*, and *GZMB*), inhibitory receptor/exhaustion programs (*PDCD1*, *CTLA4*, *ENTPD1*, *HAVCR2*, *LAG3* and *TIGIT*), tissue residency (*ITGAE*), dysfunctional/exhausted differentiation (*CXCL13*) and proliferation (*MKI67*). Colors indicate normalized gene expression levels (GEX).

(C,D) Validation and characterization of HCC-derived epitopes. Single HLA-I expressing 293T cells were pulsed with the indicated peptides at varying concentrations for 2 h (A) or at 50 μM (B), washed, and co-incubated with TCR-transduced reporter T cells for 24 h. Peptide titration assays were used to determine EC_50_ values (A), whereas peptide truncation analysis defined the minimal epitope length supporting maximal TCR recognition (B). Experiments are group per HLA class (HLA-A*02:01, HLA-A*03:01 and HLA-C*06:02) and the tested TCR is indicated above each experiment (e.g., T02 and T03 for HLA-A*02:01). T-cell activation was assessed by measuring GFP expression using flow cytometry. Measurements were performed in duplicate. Error bars represent the 95% confidence interval.

Figure S8. Characterization of tumor antigens identified in the hepatocellular carcinoma patient

(A,B) Gene expression profiles of the antigen encoding sequences from deorphanized tumor-infiltrating lymphocytes (TILs) in normal tissues from the Genotype-Tissue Expression (GTEx, blue) portal and tumors from The Cancer Genome Atlas (TCGA, orange). For antigens represented by annotated genes, expression is shown in transcripts per million (TPM; log scale) (B). For antigens encoded by transcripts not represented in GENCODE v26, expression was quantified at the genomic locus level using normalized RNAseq read coverage (C). Each dot represents an individual sample. Numbers and colors above each tissue or cancer type indicate the percentage of samples with expression exceeding the 95th percentile of normal tissue expression (P_95_), excluding testis. Cancer types are abbreviated according to TCGA nomenclature. LIHC stands for the TCGA Liver Hepatocellular Carcinoma dataset.

(C) UMAP visualizing *ENSG00000297530 and SOAT2* expression in single-cell data of pre- and post-treatment biopsies from the hepatocellular carcinoma patient. Cells are colored according to normalized *ENSG00000297530 and SOAT2* expression density. Major cell populations are indicated on the UMAP, including cancer cells, T cells, B cells, myeloid cells, endothelial cells, pDCs, and fibroblasts.

(D,E) Epitope mapping and validation of DCAF4L2 HLA-A*02:01 restricted response. DCAF4L2 truncation constructs were generated to identify the coding region encoding the epitope recognized by HLA-A*02:01 restricted patient-derived TCRs. The *DCAF4L2* transcript is shown schematically in (D), with non-recognized coding-sequence subsequences shown in gray and recognized regions in green. The HLA-C*06:02 epitopes and the distal HLA-A*02:01 epitope-encoding region are indicated in grey and green respectively. In (E), HLA-A*02:01 293T cells expressing full-length DCAF4L2, the indicated truncation constructs (Trim 1–6), or no antigen control were assessed for recognition by the indicated DCAF4L2-specific TCRs (T417 and T612) expressed in SL8 reporter T cells. T cell activation was assessed by measuring GFP expression using flow cytometry. Green histograms indicate recognition of constructs containing the corresponding epitope-encoding region.

(F) Functional validation of the DCAF4L2-derived epitope in the DCAF4L2+ HCC cell line PLC/PRF/5. Target cells were modified to express HLA-C*06:02 and RFP and co-cultured with healthy donor PBMCs transduced with TCRs (either T07 or T19) recognizing the DCAF4L2-derived epitope or untransduced control PBMCs.

(G) Longitudinal quantification of images shown in figure 6G. Mean values are shown, error bars represent the 95% confidence interval.

## STAR★METHODS

### KEY RESOURCES TABLE

| REAGENT or RESOURCE | SOURCE | IDENTIFIER |
| --- | --- | --- |
| **Antibodies** |  |  |
| Ultra-LEAF(TM) Purified anti-human CD3 | Biolegend | Cat# 317326,<br>RRID:AB_11150592 |
| APC anti-human CD3 | Biolegend | Cat# 317317,<br>RRID:AB_1937213 |
| BV421 anti-human TCR $\alpha/\beta$ | Biolegend | Cat# 306721,<br>RRID:AB_2562077 |
| APC anti-mouse TCR $\beta$ chain | Biolegend | Cat# 109212,<br>RRID:AB_313435 |
| FITC anti-human HLA-A,B,C | Biolegend | Cat# 311403,<br>RRID:AB_314872 |
| FITC anti-human CD8a | Biolegend | Cat# 301006,<br>RRID:AB_314124 |
| BV421 anti-human TCR $\alpha/\beta$ | Biolegend | Cat# 306721,<br>RRID:AB_2562077 |
| FITC anti-human HLA-A,B,C | Biolegend | Cat# 311403,<br>RRID:AB_314872 |
| **Bacterial and virus strains** |  |  |
| One Shot™ OmniMAX™ 2 T1<sup>R</sup> Chemically Competent *E. coli* | Thermo Fisher Scientific | Cat# C854003 |
| Endura Electrocompetent Cells | Lucigen | Cat# 60242-1 |
| **Biological samples** |  |  |
| Healthy donor buffy coat | Belgian Red Cross-Flanders Blood Service | N/A |
| Chemicals, peptides, and recombinant proteins |  |  |
| Human IL-2 IS, premium grade | Miltenyi | 130-097-744 |
| Recombinant Human IL-7 Protein, Animal-Free | Bio-Techne | BT-007-AFL-025 |
| Human IL-15, Animal-Free Recombinant Protein | Thermo Fisher Scientific | AF-200-15-10UG |
| PepPower Chemical Peptide Synthesis, Purity $\geq 85\%$ | GenScript | SC1208 |
| Critical commercial assays |  |  |
| NEBNext Ultra II Directional RNA Library Prep Ki | New England Biolabs | E7760S |
| NEBuilder HiFi DNA Assembly Master Mix | New England Biolabs | E2621L |
| GeneJET Plasmid Miniprep Kit | Thermo Fisher Scientific | K0503 |
| Platinum Direct PCR Universal Master Mix | Thermo Fisher Scientific | A44647100 |
| **Deposited data** |  |  |
| Single-cell RNA-seq and TCR-seq | European Genome-phenome Archive | EGAS00001004809 |
| Single-cell RNA-seq and TCR-seq | National Center for Biotechnology Information | PRJNA985415 |
| **Experimental models: Cell lines** |  |  |
| HEK293T | ATCC | Cat# CRL-3216,<br>RRID:CVCL_0063 |
| Jurkat E6.1 | ATCC | Cat# TIB-152, RRID:CVCL_0367 |
| SH-SY5Y | ATCC | Cat# CRL-2266, RRID:CVCL_0019 |
| PLC/PRF/5 | ATCC | Cat# CRL-8024, RRID:CVCL_0485 |
| SNU-475 | ATCC | Cat# CRL-2236, RRID:CVCL_0497 |
| Plat-GP | Cell Biolabs | Cat# RV-103, RRID:CVCL_B490 |
| 2D3 | Campillo-Davo, D.; Versteven, M.; Roex, G.; De Reu, H.; van der Heijden, S.; Anguille, S.; Berneman, Z.N.; Van Tendeloo, V.F.I.; Lion, E. Rapid Assessment of Functional Avidity of Tumor-Specific T Cell Receptors Using an Antigen-Presenting Tumor Cell Line Electroporated with Full-Length Tumor Antigen mRNA. *Cancers* **2020**, *12*, 256. https://doi.org/10.3390/cancers12020256 |  |
| SL8 | This paper |  |
| 293T HLA-I KO | This paper |  |
| **Oligonucleotides** |  |  |
| See Table S3 |  |  |
| **Recombinant DNA** |  |  |
| eBlocks | Integrated DNA Technologies |  |
| pSIRV-NFAT-eGFP | Addgene | RRID:Addgene_118031 |
| 8xNFAT-ZsG-hCD8 | Addgene | RRID:Addgene_153417 |
| pMP71-Kaede | Addgene | RRID:Addgene_141352 |
| pMXs-GFP | Cell Biolabs | Cat# RTV-050 |
| MSGV Mouse Acceptor PGK NGFR | Addgene | RRID:Addgene_128482 |
| murine TCR OTII-2A.pMIG II | Addgene | RRID:Addgene_52112 |
| pWPXL | Addgene | RRID:Addgene_12257 |
| pWPT-GFP | Addgene | RRID:Addgene_12255 |
| pLVX-EF1alpha-2xGFP:NES-IRES-2xRFP-NLS | Addgene | RRID:Addgene_71396 |
| pLenti6.2 mTagBFP2 | Addgene | RRID:Addgene_113725 |
| pSpCas9(BB)-2A-GFP (PX458) | Addgene | RRID:Addgene_48138 |
| **Software and algorithms** |  |  |
| Cell Ranger | 10x Genomics<br>Zheng, G. X. Y. et al. (2017). Massively parallel digital transcriptional profiling of single cells. *Nature Communications* **8**: 1-12, [doi:10.1038/ncomms14049](https://doi.org/10.1038/ncomms14049) |  |
| Seurat | Stuart T, Butler A, Hoffman P, Hafemeister C, Papalexi E, III WMM, Hao Y, Stoeckius M, Smibert P, Satija R (2019). "Comprehensive Integration of Single-Cell Data." *Cell*, **177**, 1888-1902. [doi:10.1016/j.cell.2019.05.031](https://doi.org/10.1016/j.cell.2019.05.031). https://doi.org/10.1016/j.cell.2019.05.031. |  |
| Recount3 | Wilks C, Zheng SC, Chen FY, Charles R, Solomon B, Ling JP, Imada EL, Zhang D, Joseph L, Leek JT, Jaffe AE, Nellore A, Collado-Torres L, Hansen KD, Langmead B (2021). "recount3: summaries and queries for large-scale RNA-seq expression and splicing." *Genome Biol.* [doi:10.1186/s13059-021-02533-6](https://doi.org/10.1186/s13059-021-02533-6). https://doi.org/10.1186/s13059-021-02533-6. |  |
| FlowJo | BD Life Sciences |  |
| Spinnaker SDK | Teledyne Vision Solutions |  |
| Zaber motion Python API | Zaber |  |
| pyfirmata2 | https://github.com/berndporr/pyFirmata2 |  |

### EXPERIMENTAL MODEL AND STUDY PARTICIPANT DETAILS

#### Cell lines

HEK293T, PlatGP, SH-SY5Y, PLC/PRF/5 and SNU-475 were cultured in DMEM with 10% (v/v) FBS (Gibco #A5256701), 100 units/mL penicillin and 100 units/ml streptomycin (Gibco #151401220). Jurkat E6.1, 2D3 and SL8 were cultured in IMDM with 10% (v/v) FBS (Gibco #A5256701), 100 units/mL penicillin and 100 units/ml streptomycin (Gibco #151401220), supplemented with MEM Non-Essential Amino Acids Solution (Gibco #11140035) and 2mM glutamine (Gibco # 25030081).

#### Primary T cells

Blood buffy coats were obtained from Belgian Red Cross-Flanders Blood Service under protocol RKOV_19015. Primary blood mononuclear cells (PBMCs) were purified on a Ficoll gradient (GE Healthcare #17-5442-02) and cultured in IMDM with 10% (v/v) FBS (Gibco #A5256701), 100 units/mL penicillin and 100 units/ml streptomycin (Gibco #151401220), supplemented with MEM Non-Essential Amino Acids Solution (Gibco #11140035), 2 mM glutamine (Gibco # 25030081) 5 ng/ml IL-2, 10 ng/ml IL-7 and 25 ng/ml IL-15 .

#### Participant details

Patient BT21, a 62-year-old male previously diagnosed with melanoma, was treated for a brain metastasis at the University Hospital Mannheim following written consent. The study was approved by the institutional review board (Ethikkommission 2019-643N) and reported previously in Tan et al.^25^

The breast cancer patient was a post-menopausal female previously diagnosed with a non-metastatic operable newly diagnosed primary invasive carcinoma of the breast that was confirmed histologically as a pT3N0M0 ER-/PR-/HER2+ breast cancer. The patient was enrolled in the BioKey study, which was approved by the local medical ethics committee of the University Hospitals Leuven (S60100). The study was conducted according to EU legislation regarding ethical regulations and was registered online as NCT03197389. The patient provided written informed consent. The study was reported previously in Bassez et al.^1^ The patient received a single dose of pembrolizumab (anti-PD1) before surgical resection 8 days later. A pre-treatment biopsy (prior to 1 dose of pembrolizumab) was collected and an on-treatment tumor resection sample were single-cell profiled, as reported in Bassez et al.^1^

The hepatocellular carcinoma patient was a 68-year-old male previously diagnosed with an advanced hepatocellular carcinoma (pT2N0M0, BCLC-B) and was treated at University Hospitals Leuven receiving standard-of-care atezolizumab (anti-PD-L1) plus bevacizumab (anti-VEGF) before surgical resection 4 months after. The study was conducted according to EU legislation regarding ethical regulations and registered as a sample acquisition study (S62548). A pre-treatment (prior to 1 dose of pembrolizumab) and an post-treatment tumor sample (collected during resection) were single-cell profiled using Chromium Single Cell V(D)J Solution from 10x Genomics according to the manufacturer’s instructions and analyzed as reported in Cappuyns et al.^44^

### METHOD DETAILS

#### Theoretical Transcript Coverage in TWISTAR cDNA Libraries

Our TWISTAR library assembly method relies on cloning cDNA fragments with a median size of ∼300 nucleotides. Given that the median transcript length is approximately 2 kb after poly(A) capture, we can estimate the theoretical transcript coverage of the library as follows. Assuming 2 million APCs per HLA allele per screening plate, with each expressing on average one cDNA fragment, the library would represent up to approximately 0.3 million transcripts: 2 × 10^6^ ÷ (2000/300). Because cloning is directional, but the translation frame is random, only one third of inserts are expected to be translated within the correct reading frame (at least when the insert does not include the endogenous translation start site). The effective screening depth corresponds 0.1 × 10^6^ transcript-equivalent fragments. Thus, under these assumptions a transcript expressed at 10 TPM would be expected to be represented approximately once in the in-frame library. Of note, increasing the number of screening plates per HLA allele increases the screening depth and thereby improves sensitivity for lowly expressed antigens.

#### T cell clonotype definition and single-cell TCR id assignment

Given that T cells can express up to two functional TRA and TRB chains, potentially resulting in up to four distinct TCRs per single cell, without being flagged as doublets in scTCR-seq data, and considering (i) the limited sensitivity of transcript detection (such that failure to detect a given chain does not imply absence of expression) and (ii) the possibility of chain misattribution due to technical or in silico artifacts, we adapted the definition of T cell clonotypes according to the type of analysis performed. For general repertoire analyses, clonotypes were defined based solely on the most highly expressed TRB CDR3 sequence per cell. This strategy prevents artificial inflation of clonotype numbers that may arise when both TRA and TRB chains are considered, which could otherwise result in a single cell being associated with up to four complete TCRs or multiple single-chain clonotypes. For analyses focused on the frequency and phenotype associated with defined TCRs (i.e., specific TRA-TRB pairs), TCR identities were assigned to single cells only when both TRA and TRB CDR3 sequences were detected. This stringent approach ensures unambiguous identification of cells expressing the TCRs of interest but excludes cells in which one chain was not detected. When up to two TRA or TRB chains were recurrently detected within the same cell, suggesting expression of two TCRs in a group of T cells, if a TCR could be deorphanized using TWISTAR, the corresponding TCR identity was assigned to the multi-TCR-expressing cell. If none could be deorphanized, the identity of the first TCR in alphabetical order was assigned to avoid dual labeling of single cells.

#### Lentivirus production and transduction

Lentiviral particles were produced by transient transfection of HEK293T cells with the lentiviral transfer vector together with plasmids encoding Tat, Rev, Gag-Pol, and VSV-G using FuGENE 4K transfection reagent (Promega). Viral supernatants were collected after 48h and used for transduction of target cells. For HEK293T-derived and tumor cell lines, cells were transduced with viral supernatant without additional transduction enhancers. For Jurkat-derived cell lines and primary T cells, transductions were performed in the presence of 8 µg/mL polybrene (Sigma) through a spinoculation at 800 × g for 45 min at 32°C. The average multiplicity of infection (MOI) was estimated from the fraction of transduced cells using the Poisson model, where MOI = −ln(1 − P), with P representing the percentage of transduced cells.

#### Generation of single-HLA-presenting cells

A guide RNA (gRNA) targeting the sequence 5′-GATGTAATCCTTGCCGTCGT-3′ was selected based on conservation across *HLA-A*, *HLA-B*, and *HLA-C* alleles expressed in HEK293T cells and the absence of predicted exonic off-target sites identified using CCTOP (https://cctop.cos.uni-heidelberg.de). It was cloned into the pX458 (also known as pSpCas9(BB)-2A-GFP) plasmid and HEK293T cells were transfected with the resulting plasmid using FuGENE HD transfection reagent (Promega) to achieve transient co-expression of Cas9 and the gRNA. Following transfection and expansion for 2 weeks, cells were stained with a FITC-conjugated anti-human HLA-A,B,C antibody and HLA-I-negative cells were single-cell sorted into 96-well plates using a BD FACSAria Fusion cell sorter. Clonal populations were expanded and screened for loss of endogenous HLA-I surface expression. A clone exhibiting complete loss of endogenous HLA-I surface expression and the highest proliferation rate was selected as HLA-I-null parental line. Individual HLA alleles were then stably expressed in the HLA-I-null cells by lentiviral transduction using an EF1α promoter-driven pWPXL vector at MOI of 1. HLA surface expression was confirmed by flow cytometry and positives cells sorted and expanded for downstream experiments.

#### Generation of the Jurkat E6.1 based NFAT-GFP reporter population

VSV-G-pseudotyped retroviral particles were produced by transient transfection of Plat-GP packaging cells (Cell Biolabs) with pCMV-VSV-G and pSIRV-NFAT-eGFP vectors. Jurkat E6.1 cells were transduced using viral supernatants collected 48h post-transfection, filtered through a 0.45-µm membrane, and pre-loaded onto Retronectin-coated plates (Takara Bio). Following transduction, cells were stimulated on anti-CD3-coated plates to induce NFAT-dependent GFP expression. Cells exhibiting the highest GFP fluorescence following stimulation were bulk sorted using a BD FACSAria Fusion cell sorter to establish the NFAT-GFP reporter population.

#### Knockout of endogenous *TRA* and *TRB* genes

To disrupt the coding sequences of the endogenous TCR expressed in Jurkat E6.1 NFAT-GFP reporter cells, gRNAs targeting the sequences 5′-CTGGGTCACCGACTGGGCTCTGG-3′ and 5′-ACTCGCTACCAGGATGCAAAGGG-3′ within *TRAV8-4* and *TRBV12-3*, respectively, were cloned into pX458. The resulting plasmids were introduced sequentially by electroporation using the Lonza Amaxa 4D-Nucleofector System and SE Cell Line 4D kit. First, two weeks after TRB targeting, cells were stained with APC-conjugated anti-human CD3 antibody and CD3-negative cells were bulk sorted. Because cell surface CD3 expression depends on the assembly of a functional TCRαβ complex, loss of CD3 expression was used as a surrogate marker for successful TRB disruption. Then, the resulting population was subsequently electroporated with TRAV8-4 targeting pX458 vector for TRA disruption. Individual cells were single -cell sorted into individual wells of a 96-well plate based on expression of the pX458-encoded GFP transfection marker using a BD FACSAria Fusion cell sorter. Clonal populations were expanded and screened for loss of endogenous TRA expression by transduction with lentiviral vectors encoding the Jurkat TRB chain, followed by assessment of CD3 surface expression.

#### Selection of an NFAT-GFP reporter clonal cell line for CD8 TCR studies

NFAT-GFP TCR knockout clonal populations (n = 48) were transduced with lentiviral vectors encoding CD8 (using the pWPXL vector carrying the CD8α-P2A-CD8β cassette derived from the 8xNFAT-ZsG-hCD8 construct) together with a TCR specific for the PIK3CA_H1047L-derived epitope ALHGGWTTK (pWPXL KC-TCR2 vector). The selected TCR (CDR3α: CALTVGGSYIPTF, CDR3β: CASSQGGQGWRETQYF) was previously reported to exhibit micromolar functional avidity and CD8 dependence (Chandran et al.^21^). Transduced populations were co-incubated for 24h with HEK293T cells stably expressing a PIK3CA_H1047L-encoding minigene together with the respective HLA allele HLA-A*03:01. T cell activation was assessed by flow cytometric quantification of GFP induction within CD3-positive cells. The clone yielding the highest proportion of GFP-positive cells was selected. To generate the final reporter line, the selected clone was transduced exclusively with the CD8-encoding lentiviral vector, bulk sorted, and designated SL8. This cell line was used for all downstream TWISTAR experiments.

#### TCR construct design and cloning

Unless otherwise specified, TCR expression constructs were assembled in a pWPXL-derived lentiviral backbone containing the TRAC constant region to generate bicistronic TRB-P2A-TRA expression cassettes. Variable regions were designed based on paired-chain contig outputs from Cell Ranger filtered contig analysis and synthesized as IDT eBlocks gene fragments to generate bicistronic TCR constructs encoding TRB-P2A-TRA. The constructs were assembled using NEBuilder HiFi DNA Assembly (New England Biolabs) with the following overlapping homology regions:

• pWPXL / TRBV: 5′-acgagactagcctcgaggtttaaactacgggatccGCCACC-3′

• TRBV / TRBC1-P2A: 5′-AGGACCTGAACAAGGTGTTCCCACCC-3′

• TRBV / TRBC2-P2A: 5′-AGGACCTGAAAAACGTGTTCCCACCC-3′

• TRBC1/2-P2A / TRAV: 5′-ACGTGGAGGAGAACCCTGGACCT-3′

• TRAV / TRAC: 5′-ATATCCAGAACCCTGACCCTGCCGTGTACC-3′

#### TWISTAR cDNA lentiviral library construct

Total RNA from tumor cell lines or tumor biopsies was isolated using TRIzol (Thermo Fisher Scientific). Eight hundred nanograms of RNA were used for poly(A)+ mRNA enrichment and cDNA-based libraries were generated based on the directional RNAseq library preparation protocol using the NEBNext Poly(A) mRNA Magnetic Isolation Module and NEBNext Ultra II Directional RNA Library Prep Kit (New England Biolabs). Libraries were prepared with a target insert size of 300 base pairs according to the manufacturer’s instructions except that following adaptor ligation, PCR amplification was performed using TCP1v2 and TCP2v2 primers (Table S3) to introduce homology regions compatible with the lentiviral antigen expression backbone. Specifically, cDNA amplicons were designed for cloning into a modified pLenti6.2-mTagBFP2 vector. The regions flanking the mTagBFP2 cassette were engineered to be compatible with the cDNA library adaptor sequences, allowing the cDNA insert to replace the CMV promoter-driven mTagBFP2 sequence upon ligation. In addition, the original puromycin resistance cassette (PuroR) was replaced with an mCherry-P2A-PuroR cassette, enabling both MOI determination by flow cytometry and visual identification of transduction by fluorescence microscopy. The vector was linearized using BsrGI and MluI digestion (releasing mTagBFP2 coding sequence). PCR products were assembled into 100 ng of linearized destination vector using NEBuilder HiFi DNA Assembly (New England Biolabs) at a 1:7 vector-to-insert molar ratio. Assembly reactions were transformed into Endura electrocompetent cells (Lucigen) and plated onto LB agar containing 100 µg/mL carbenicillin using eight 245 × 245 mm square dishes. Plates were incubated overnight at 32°C. Transformants (about 10 million in total) were then harvested, pooled, and plasmid DNA was extracted from one-tenth of the total bacterial biomass.

#### TWISTAR screening setup and antigen coding sequence recovery

Each patient expressed up to 6 different HLA-I alleles. When conducting TWISTAR experiments, we generated single HLA expressing HEK293T cells. This was done at a multiplicity of infection (MOI) of 1 with the autologous tumor derived cDNA lentiviral library. Subsequently, transduced cells were seeded into two single-well SBS-format culture plates (VWR #734-2966), in which three HLA alleles were pooled per plate (2.10^6^ cells per condition). SL8 reporter cells (10^7^), transduced with a TCR of interest, were added to each plate, and co-cultures were incubated for 48 h at 37°C. Following incubation, entire plate surfaces were imaged using a Nucleus MVR inverted fluorescence microscope (Zaber) equipped with a 2.5x large-field objective (Zeiss #420120-9901-000) and a 1.2’’ sensor camera (FLIR BFS-U3-244S8M-C). A custom Python script (Zenodo: [DOI]) was used to identify GFP-positive clusters, and their spatial coordinates were subsequently used to guide retrieval of up to 12 positive clusters using a custom-built cell picker.

The cell picker is custom-build and consists of a blunt-end 25G stainless steel needle (Metcal #925050-TE) connected via a 1 mm inner-diameter polyethylene tubing (Braun #8255059) to a 1 mL Luer-Lok syringe (BD #309628), mounted on an Arduino-controlled motorized positioning system (MGN12H linear guides, 8 mm lead screw, NEMA17 stepper motors; assembly files available upon request). Cells were aspirated in TrypLE to ensure dissociation and maintain viability.

Picked cells were transferred into 96-well plates pre-coated with 5 µg/cm² collagen I (Thermo Fisher Scientific #A1048301) and expanded. Confirmation of the immunogenic signal is performed 3 days later by adding 10^4^ cognate TCR-expressing SL8 reporter cells per well, followed by imaging the next day. Confirmed antigen-reactive populations were isolated again, and antigen-encoding sequences were amplified by PCR using Platinum Direct PCR Universal Master Mix (Thermo Fisher Scientific) with TRP1 and TRP2 primers (Table S3). Amplicons were cloned into pLenti6.2 vector and functionally validated for their ability to trigger recognition by TCR-expressing SL8 cells. Sequence identity was then determined by Sanger sequencing.

#### Minigenes recognition assays

For experiments where potential GFP+ clusters were validated, for experiments where antigen-coding sequences were trimmed to identify which part of the sequence gives rise to the presented epitope, or for experiments where the exact HLA that presents the epitope is determined, 10^4^ HEK293T cells expressing a single HLA allele were seeded per well in 384-well plates. Cells were transfected with antigen-encoding pLenti6.2 vectors using FuGENE 4K (Promega) and immediately co-cultured with SL8 TCR-expressing reporter cells. GFP expression was assessed by fluorescence imaging the following day.

#### Epitope validation using peptide-pulse experiments

Following validation of TWISTAR hits and assignment of HLA restriction, peptide-HLA binding predictions were performed using NetMHCpan-4.1^45^. Peptides with predicted binding rank <2 were prioritized. Where multiple candidate peptides were identified, the antigen-encoding sequence was subdivided into overlapping fragments, which were cloned and assessed for recognition by the cognate TCR. Once an epitope could be reliably predicted *in silico*, the corresponding peptide was synthesized and tested in a peptide-pulse experiment. In 96-well plates, 6.10^4^ HEK293T cells expressing a single HLA allele were incubated with peptide concentrations ranging from 1 pM to 500 µM in IMDM supplemented with 1% FBS for 2 h at 37°C. Cells were then washed twice and co-cultured for 24h with 6.10^3^ TCR-expressing SL8 cells. EGFP induction was quantified by flow cytometry within the CD3-positive population.

#### TCR-mediated lytic activity

Primary T cells isolated from healthy blood donors were activated with 1:100 diluted T Cell TransAct reagent (Miltenyi #130-111-160). Two days later, cells were transduced with TCR-encoding lentiviral vectors in which human TCR constant regions were replaced with murine constant regions to minimize mispairing with endogenous TCR chains. Three days post-transduction, TCR-positive cells were bulk sorted using a BD FACSAria Fusion cell sorter based on mouse TCRβ surface staining. After a 5-day resting period, sorted T cells were co-cultured in 384-well plates with H2B-mScarlet2-expressing tumor cell lines or HEK293T cells expressing a single HLA allele together with the relevant tumor antigen at an effector-to-target (E:T) ratio of 10:1. Co-cultures were performed in the presence of either 1:1000 Caspase-3/7 Green dye or 1:200 Annexin V Green dye (Sartorius #4440 and #4644). Plates were imaged every 2h for 48h using an Incucyte S3 live-cell imaging system, and full-well fluorescence imaging was additionally performed daily using the MVR microscope platform. Target cell killing was quantified by monitoring reduction of the H2B-mScarlet2 fluorescent signal, while apoptosis-associated fluorescence was used for representative imaging.

#### Antigen coding transcript expression analysis

When relevant transcript annotations were available in the GENCODE v26 human reference annotation, gene expression levels were assessed using the Monorail pipeline for in-house-generated bulk RNAseq datasets (BT21 tumor cell line and tumor biopsies) or obtained directly from the recount3 resource for TCGA, GTEx, and CCLE samples^46^. When reference transcript annotations did not adequately represent the detected antigen-coding transcript, expression was evaluated using a genomic locus–based alignment depth analysis. Briefly, average per-base coverage was computed from Monorail-generated BigWig files and normalized to the total number of mapped bases for each sample, scaled to 10 billion mapped bases (normdepth).

#### Survival analysis in immunotherapy-treated HCC cohorts

Bulk RNAseq data from the GO30140 and IMbrave150 cohorts (EGAS00001005503) were used to assess the association between baseline tumor *DCAF4L2* expression and progression-free survival in immunotherapy or sorafenib treated hepatocellular carcinoma. Analyses were restricted to pre-treatment tumor samples from patients receiving atezolizumab-based therapies, including atezolizumab monotherapy or atezolizumab plus bevacizumab. *DCAF4L2* expression was dichotomized based on raw RNAseq counts (0 versus >0). Progression-free survival was estimated using the Kaplan-Meier method and compared using the log-rank test. Hazard ratios (HRs) and 95% confidence intervals were estimated using univariable Cox proportional hazards models.

### QUANTIFICATION AND STATISTICAL ANALYSIS

Statistical details of experiments can be found in the figure legends. Data analysis was performed in R. All error bars in figures indicate 95% confidence intervals.

## REFERENCES

1. Bassez, A., et al., A single-cell map of intratumoral changes during anti-PD1 treatment of patients with breast cancer. Nat Med, 2021. 27(5): p. 820–832.

2. Franken, A., et al., CD4(+) T cell activation distinguishes response to anti-PD-L1+anti-CTLA4 therapy from anti-PD-L1 monotherapy. Immunity, 2024. 57(3): p. 541–558 e7.

3. Guo, X., et al., Global characterization of T cells in non-small-cell lung cancer by single-cell sequencing. Nat Med, 2018. 24(7): p. 978–985.

4. Miller, B.C., et al., Subsets of exhausted CD8(+) T cells differentially mediate tumor control and respond to checkpoint blockade. Nat Immunol, 2019. 20(3): p. 326–336.

5. Sahin, U., et al., Individualized mRNA vaccines evoke durable T cell immunity in adjuvant TNBC. Nature, 2026. 651(8107): p. 1088–1096.

6. Weber, J.S., et al., Individualised neoantigen therapy mRNA-4157 (V940) plus pembrolizumab versus pembrolizumab monotherapy in resected melanoma (KEYNOTE-942): a randomised, phase 2b study. Lancet, 2024. 403(10427): p. 632–644.

7. Rojas, L.A., et al., Personalized RNA neoantigen vaccines stimulate T cells in pancreatic cancer. Nature, 2023. 618(7963): p. 144–150.

8. Brochier, W., O. Bricard, and P.G. Coulie, Facts and Hopes in Cancer Antigens Recognized by T Cells. Clin Cancer Res, 2023. 29(2): p. 309–315.

9. Erhard, F., et al., Identification of the Cryptic HLA-I Immunopeptidome. Cancer Immunol Res, 2020. 8(8): p. 1018–1026.

10. Cai, Y., et al., Immunopeptidomics-guided discovery and characterization of neoantigens for personalized cancer immunotherapy. Sci Adv, 2025. 11(21): p. eadv6445.

11. Chong, C., et al., Integrated proteogenomic deep sequencing and analytics accurately identify non-canonical peptides in tumor immunopeptidomes. Nat Commun, 2020. 11(1): p. 1293.

12. Joglekar, A.V. and G. Li, T cell antigen discovery. Nat Methods, 2021. 18(8): p. 873–880.

13. Meyer, M., et al., MediMer: a versatile do-it-yourself peptide-receptive MHC class I multimer platform for tumor neoantigen-specific T cell detection. Front Immunol, 2023. 14: p. 1294565.

14. Kovaleva, V.A., et al., copepodTCR: Identification of Antigen-Specific T Cell Receptors with combinatorial peptide pooling. bioRxiv, 2024.

15. Kula, T., et al., T-Scan: A Genome-wide Method for the Systematic Discovery of T Cell Epitopes. Cell, 2019. 178(4): p. 1016–1028 e13.

16. Kohlgruber, A.C., et al., High-throughput discovery of MHC class I- and II-restricted T cell epitopes using synthetic cellular circuits. Nat Biotechnol, 2025. 43(4): p. 623–634.

17. Gee, M.H., et al., Antigen Identification for Orphan T Cell Receptors Expressed on Tumor-Infiltrating Lymphocytes. Cell, 2018. 172(3): p. 549–563 e16.

18. Levi, S.T., et al., Neoantigen Identification and Response to Adoptive Cell Transfer in Anti-PD-1 Naive and Experienced Patients with Metastatic Melanoma. Clin Cancer Res, 2022. 28(14): p. 3042–3052.

19. Hu, Z., et al., Personal neoantigen vaccines induce persistent memory T cell responses and epitope spreading in patients with melanoma. Nat Med, 2021. 27(3): p. 515–525.

20. Hooijberg, E., et al., NFAT-controlled expression of GFP permits visualization and isolation of antigen-stimulated primary human T cells. Blood, 2000. 96(2): p. 459–66.

21. Chandran, S.S., et al., Immunogenicity and therapeutic targeting of a public neoantigen derived from mutated PIK3CA. Nat Med, 2022. 28(5): p. 946–957.

22. Morimoto, S., et al., Establishment of a novel platform cell line for efficient and precise evaluation of T cell receptor functional avidity. Oncotarget, 2018. 9(75): p. 34132–34141.

23. Rosskopf, S., et al., A Jurkat 76 based triple parameter reporter system to evaluate TCR functions and adoptive T cell strategies. Oncotarget, 2018. 9(25): p. 17608–17619.

24. Neefjes, J., et al., Towards a systems understanding of MHC class I and MHC class II antigen presentation. Nat Rev Immunol, 2011. 11(12): p. 823–36.

25. Tan, C.L., et al., Prediction of tumor-reactive T cell receptors from scRNA-seq data for personalized T cell therapy. Nat Biotechnol, 2025. 43(1): p. 134–142.

26. Vita, R., et al., The Immune Epitope Database (IEDB): 2018 update. Nucleic Acids Res, 2019. 47(D1): p. D339–D343.

27. Yang, L., et al., LINC00221 silencing prevents the progression of hepatocellular carcinoma through let-7a-5p-targeted inhibition of MMP11. Cancer Cell Int, 2021. 21(1): p. 202.

28. Hoffmann, M.M. and J.E. Slansky, T-cell receptor affinity in the age of cancer immunotherapy. Mol Carcinog, 2020. 59(7): p. 862–870.

29. Zhao, Q., et al., Differential evolution of MAGE genes based on expression pattern and selection pressure. PLoS One, 2012. 7(10): p. e48240.

30. Morgan, R.A., et al., Cancer regression and neurological toxicity following anti-MAGE-A3 TCR gene therapy. J Immunother, 2013. 36(2): p. 133–51.

31. Vizoso, M., et al., Epigenetic activation of a cryptic TBC1D16 transcript enhances melanoma progression by targeting EGFR. Nat Med, 2015. 21(7): p. 741–50.

32. Akavia, U.D., et al., An integrated approach to uncover drivers of cancer. Cell, 2010. 143(6): p. 1005–17.

33. Tsai, J.W., et al., FOXR2 Is an Epigenetically Regulated Pan-Cancer Oncogene That Activates ETS Transcriptional Circuits. Cancer Res, 2022. 82(17): p. 2980–3001.

34. Schmitt-Hoffner, F., et al., FOXR2 Stabilizes MYCN Protein and Identifies Non-MYCN-Amplified Neuroblastoma Patients With Unfavorable Outcome. J Clin Oncol, 2021. 39(29): p. 3217–3228.

35. Simoni, Y., et al., Bystander CD8(+) T cells are abundant and phenotypically distinct in human tumour infiltrates. Nature, 2018. 557(7706): p. 575–579.

36. Loriot, A., T. Boon, and C. De Smet, Five new human cancer-germline genes identified among 12 genes expressed in spermatogonia. Int J Cancer, 2003. 105(3): p. 371–6.

37. Wang, H., et al., DCAF4L2 promotes colorectal cancer invasion and metastasis via mediating degradation of NFkappab negative regulator PPM1B. Am J Transl Res, 2016. 8(2): p. 405–18.

38. Finn, R.S., et al., Atezolizumab plus Bevacizumab in Unresectable Hepatocellular Carcinoma. N Engl J Med, 2020. 382(20): p. 1894–1905.

39. Sethna, Z., et al., RNA neoantigen vaccines prime long-lived CD8(+) T cells in pancreatic cancer. Nature, 2025. 639(8056): p. 1042–1051.

40. Borgers, J.S.W., et al., Personalized, autologous neoantigen-specific T cell therapy in metastatic melanoma: a phase 1 trial. Nat Med, 2025. 31(3): p. 881–893.

41. Apavaloaei, A., et al., Tumor antigens preferentially derive from unmutated genomic sequences in melanoma and non-small cell lung cancer. Nat Cancer, 2025. 6(8): p. 1419–1437.

42. Van Tongelen, A., A. Loriot, and C. De Smet, Oncogenic roles of DNA hypomethylation through the activation of cancer-germline genes. Cancer Lett, 2017. 396: p. 130–137.

43. Kim, R., P. Kulkarni, and S. Hannenhalli, Derepression of Cancer/testis antigens in cancer is associated with distinct patterns of DNA hypomethylation. BMC Cancer, 2013. 13: p. 144.

44. Cappuyns, S., et al., Single-cell RNA sequencing-derived signatures define response patterns to atezolizumab + bevacizumab in advanced hepatocellular carcinoma. J Hepatol, 2025. 82(6): p. 1036–1049.

45. Reynisson, B., et al., NetMHCpan-4.1 and NetMHCIIpan-4.0: improved predictions of MHC antigen presentation by concurrent motif deconvolution and integration of MS MHC eluted ligand data. Nucleic Acids Res, 2020. 48(W1): p. W449–W454.

46. Wilks, C., et al., recount3: summaries and queries for large-scale RNA-seq expression and splicing. Genome Biol, 2021. 22(1): p. 323.

