## Supplementary figures and images for "Functional Deorphanization of Tumor-reactive TCRs Uncovers a Landscape of Shared, Immune Dominant Antigens across Cancers"

### Figures_HD

Figure 1

A

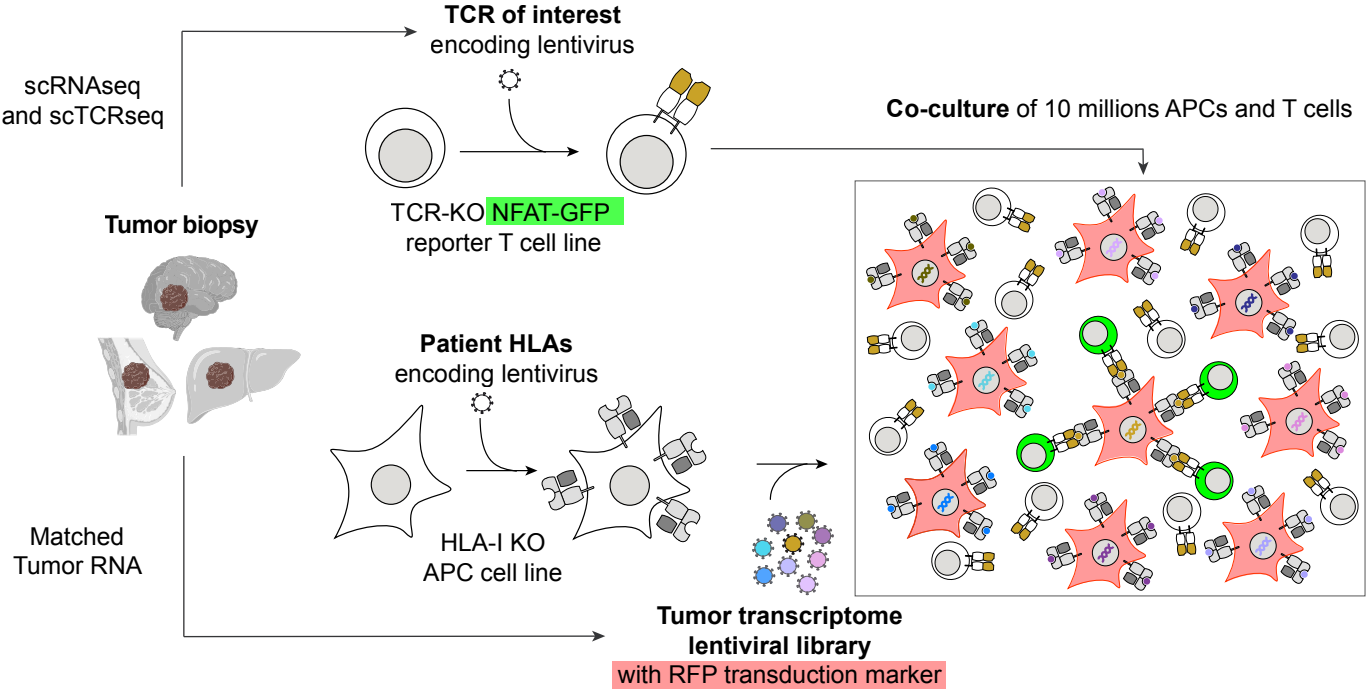

B

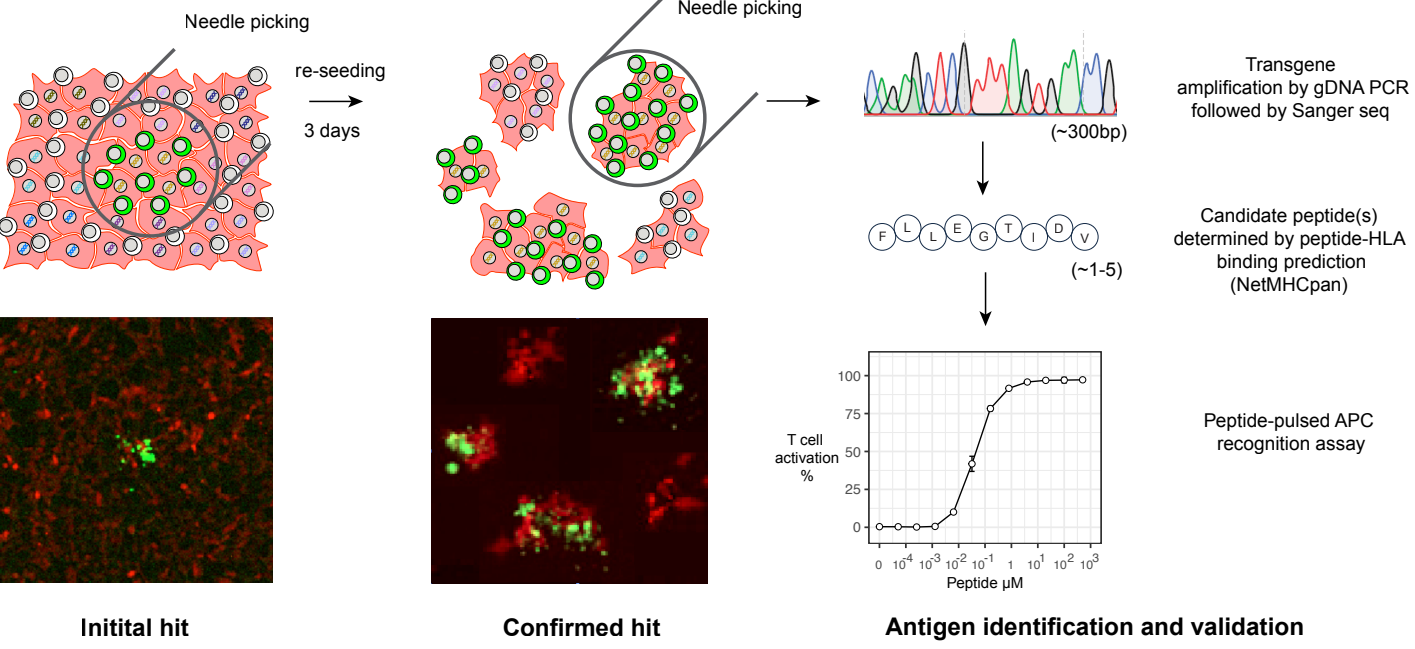

Figure 2

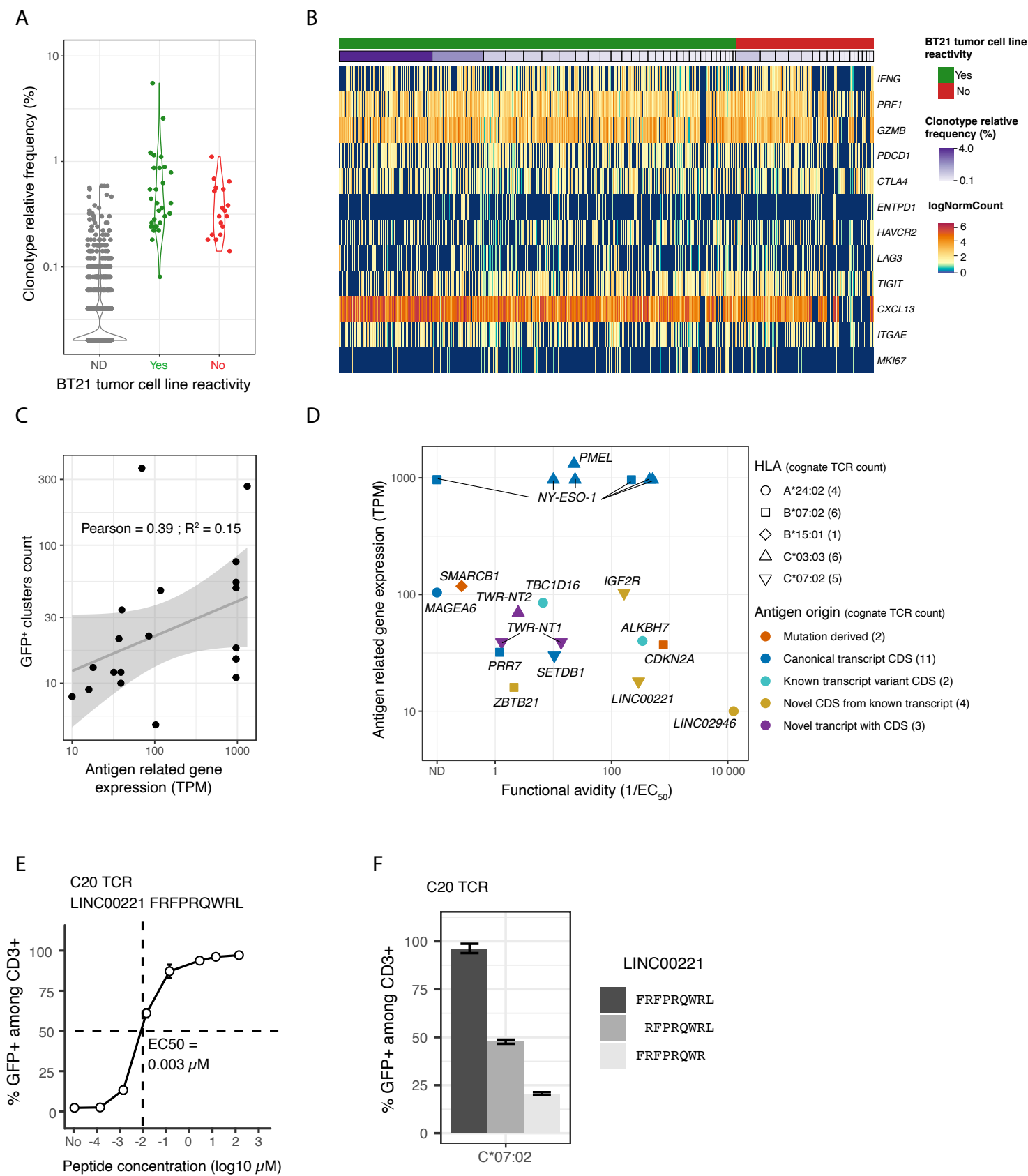

Figure 3

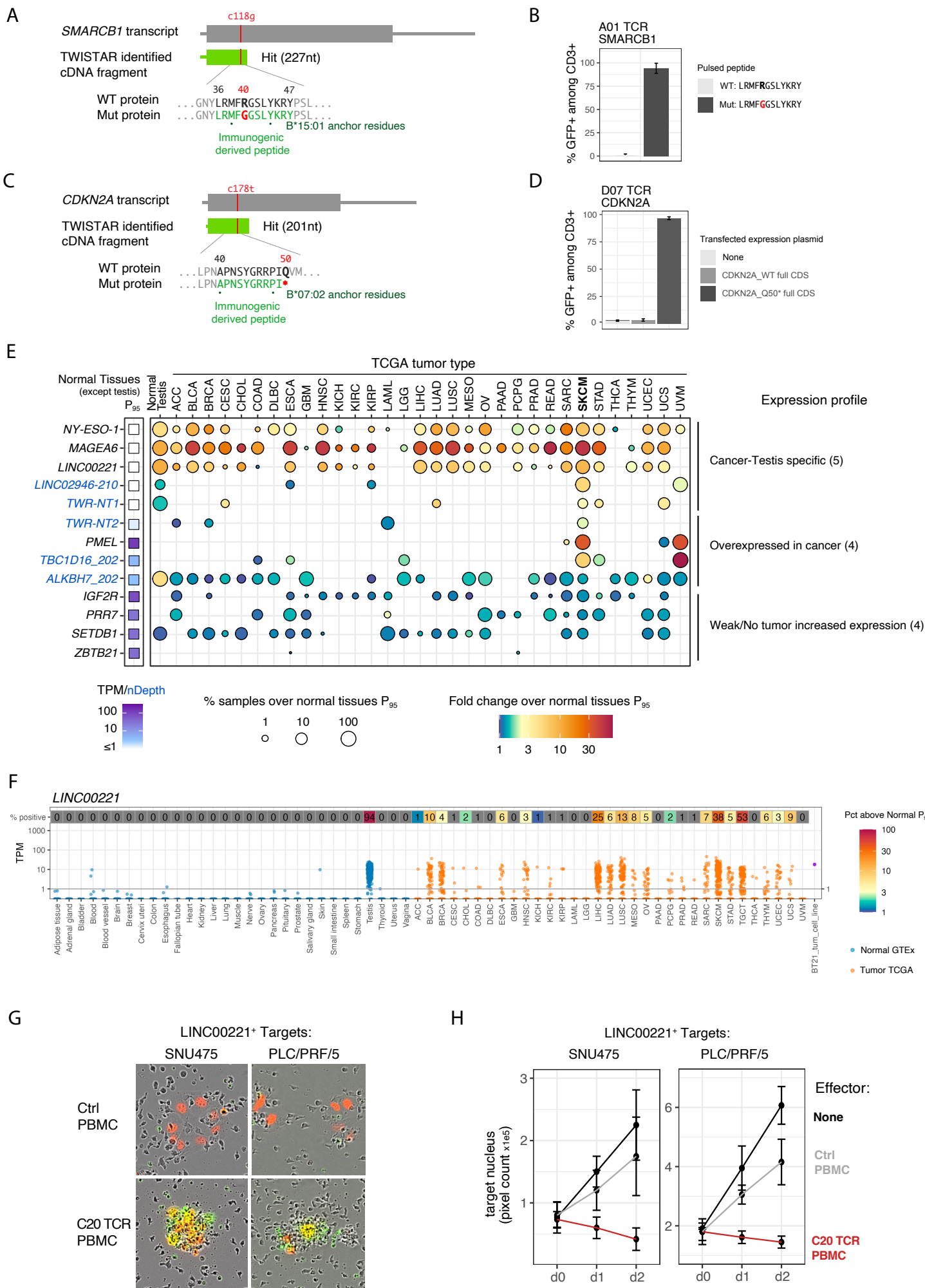

Figure 4

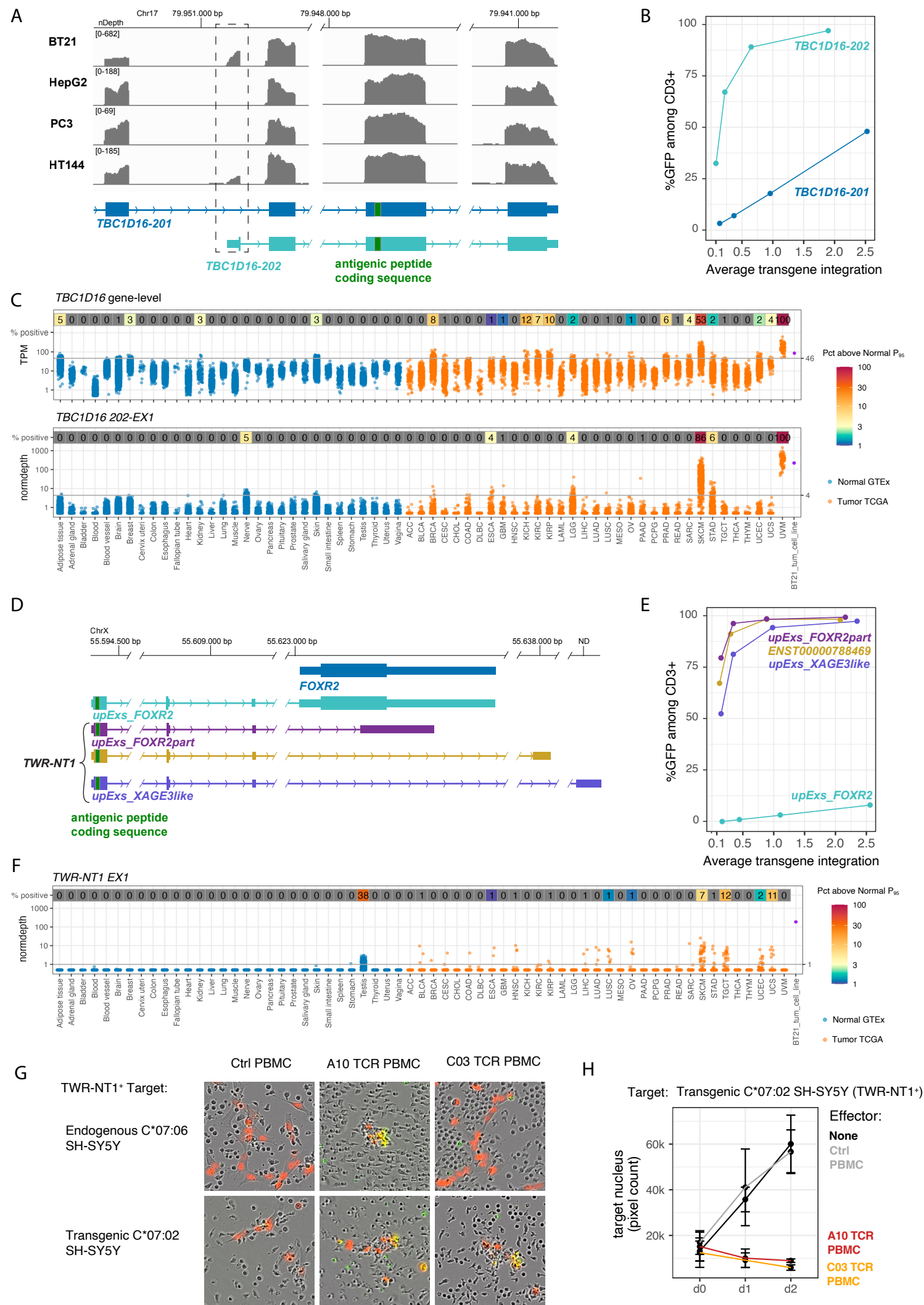

Figure 5

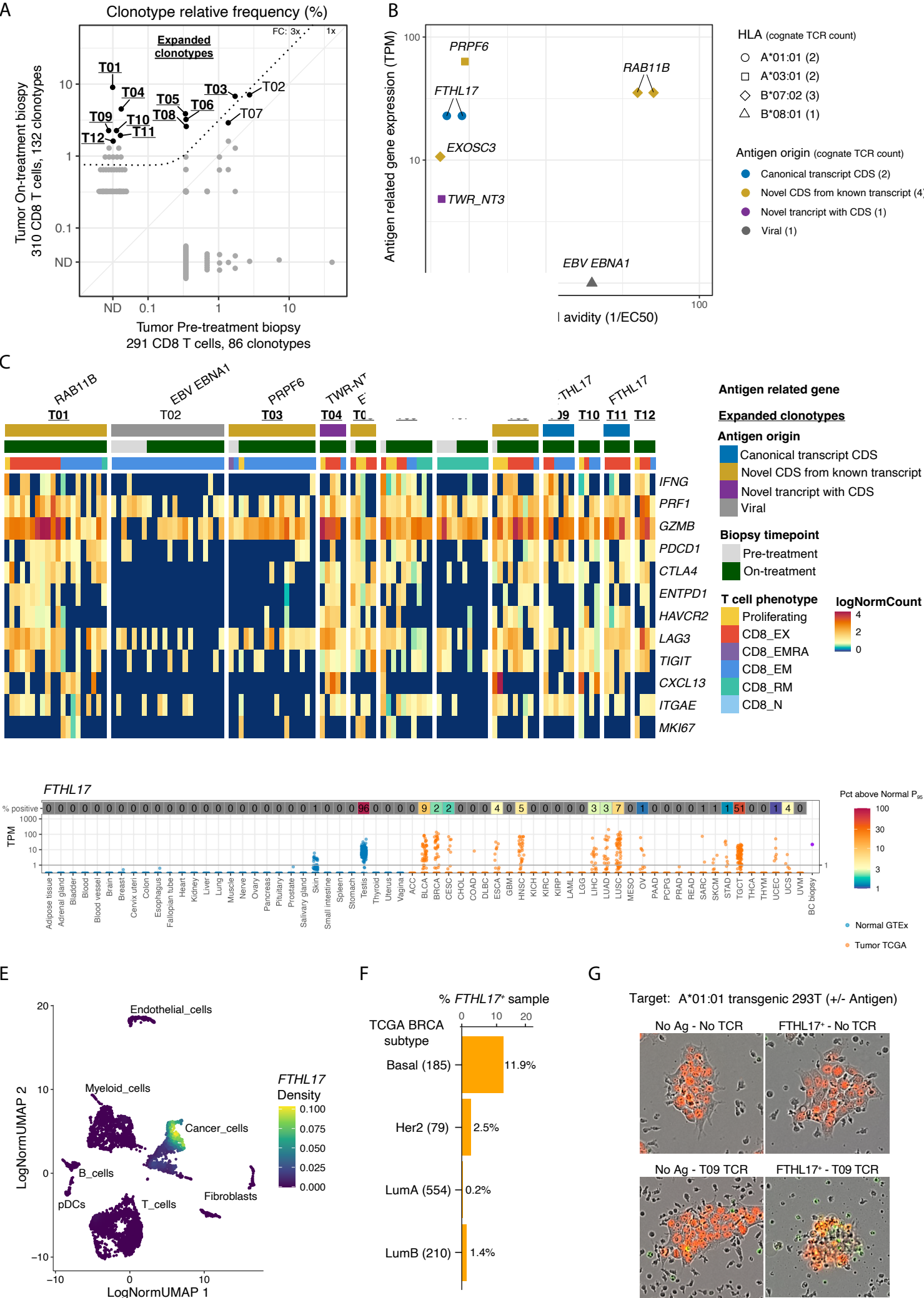

Figure 6

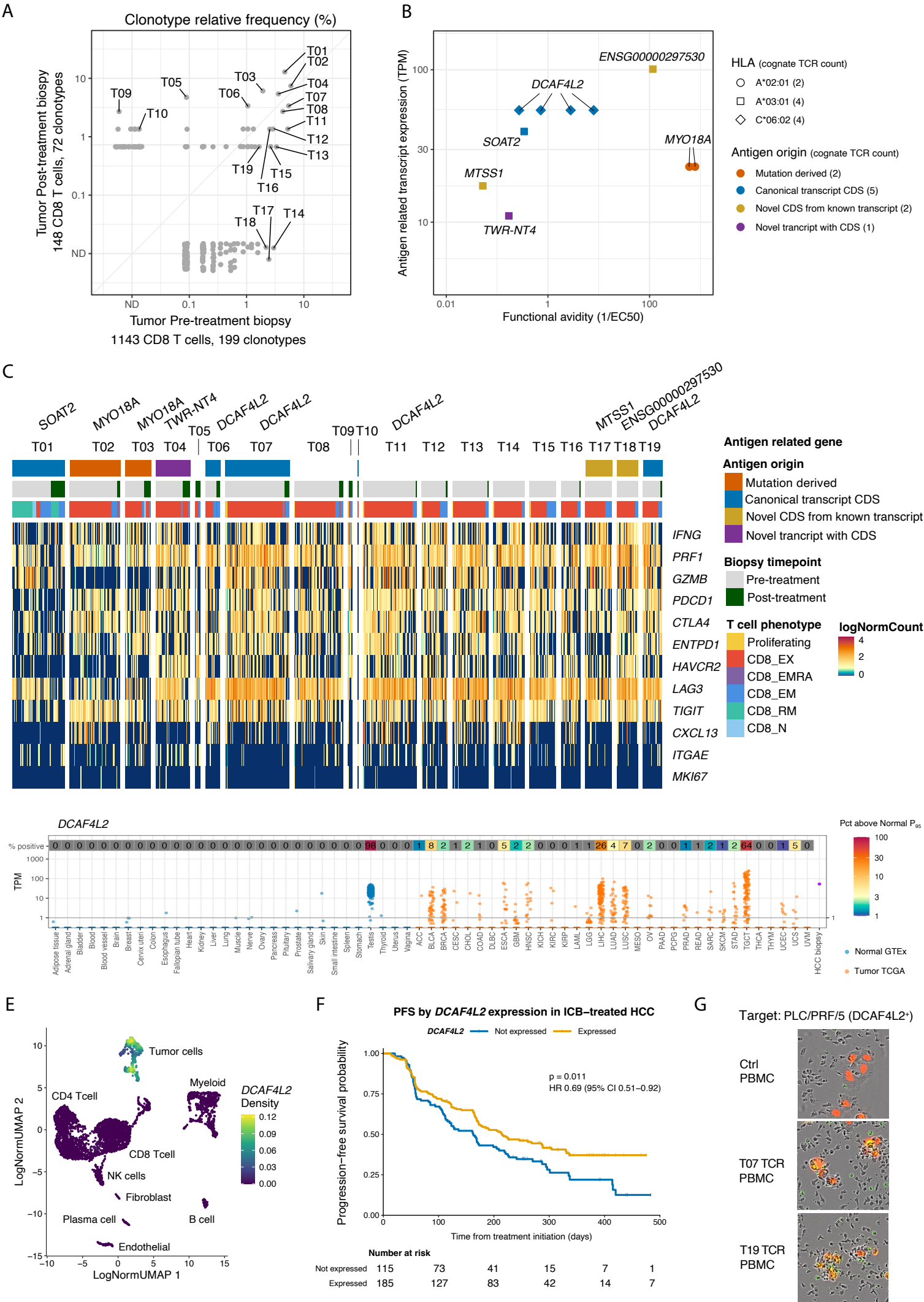
