## Supplemental Figures for "Functional Deorphanization of Tumor-reactive TCRs Uncovers a Landscape of Shared, Immune Dominant Antigens across Cancers"

Supplement figure 1

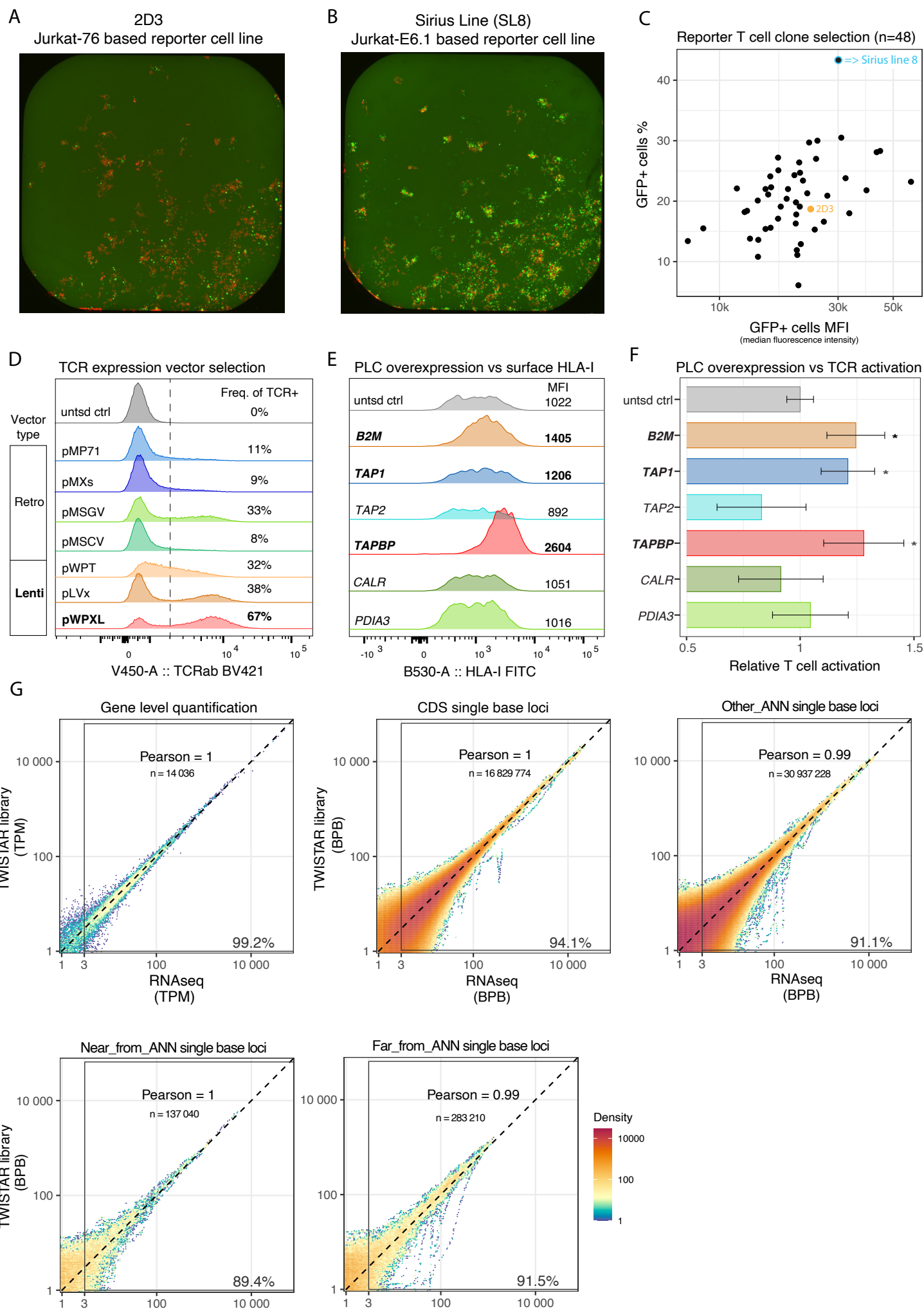

Supplement figure 2

A

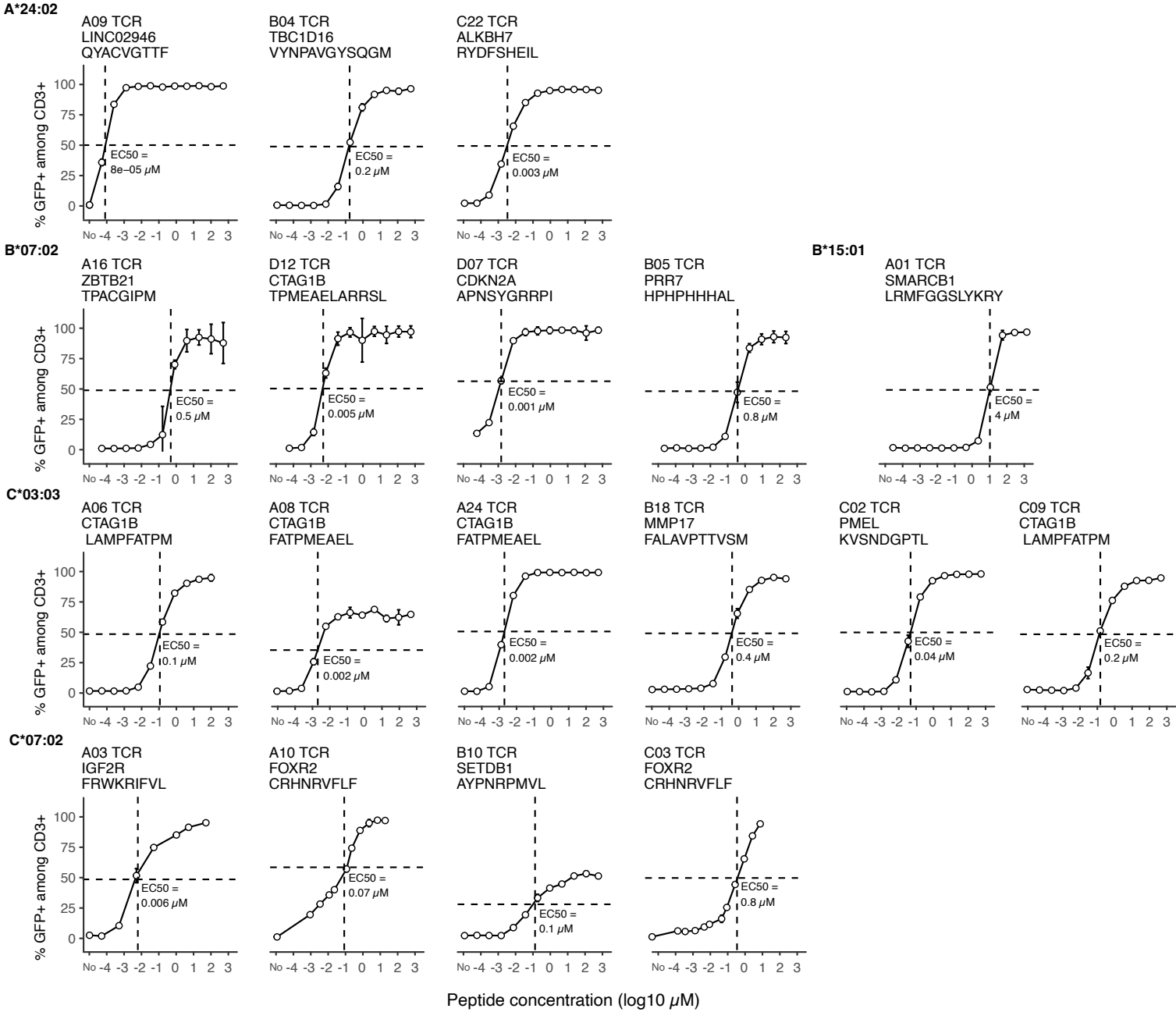

B

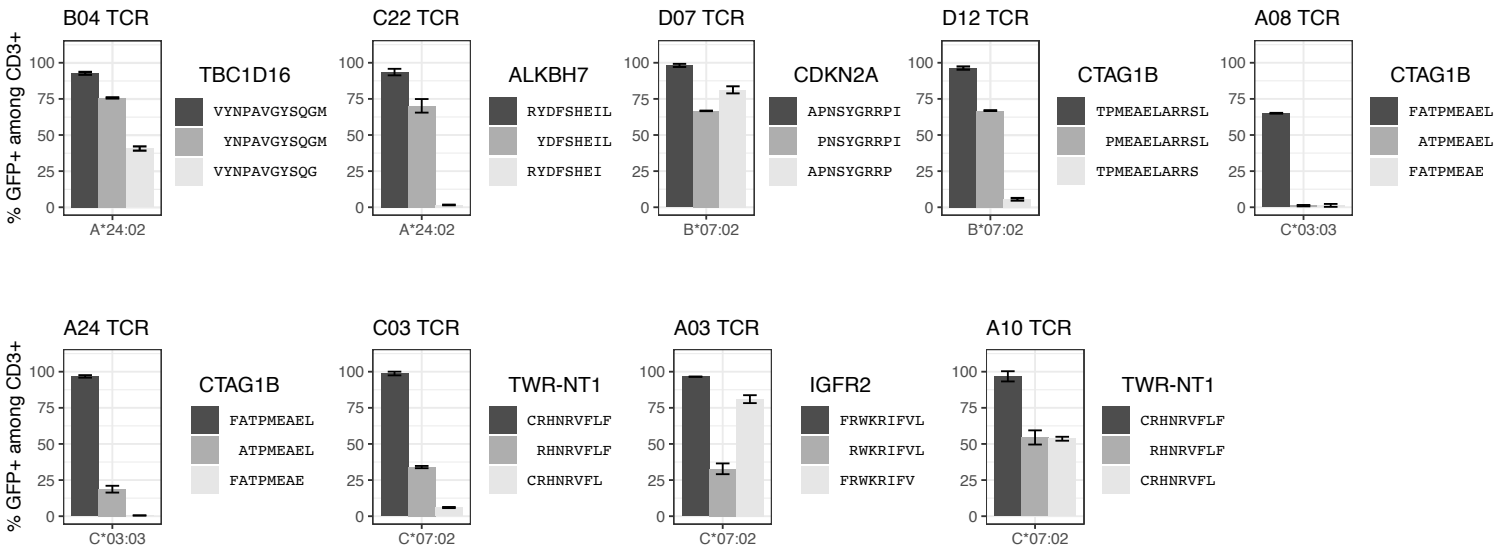

Supplement figure 3

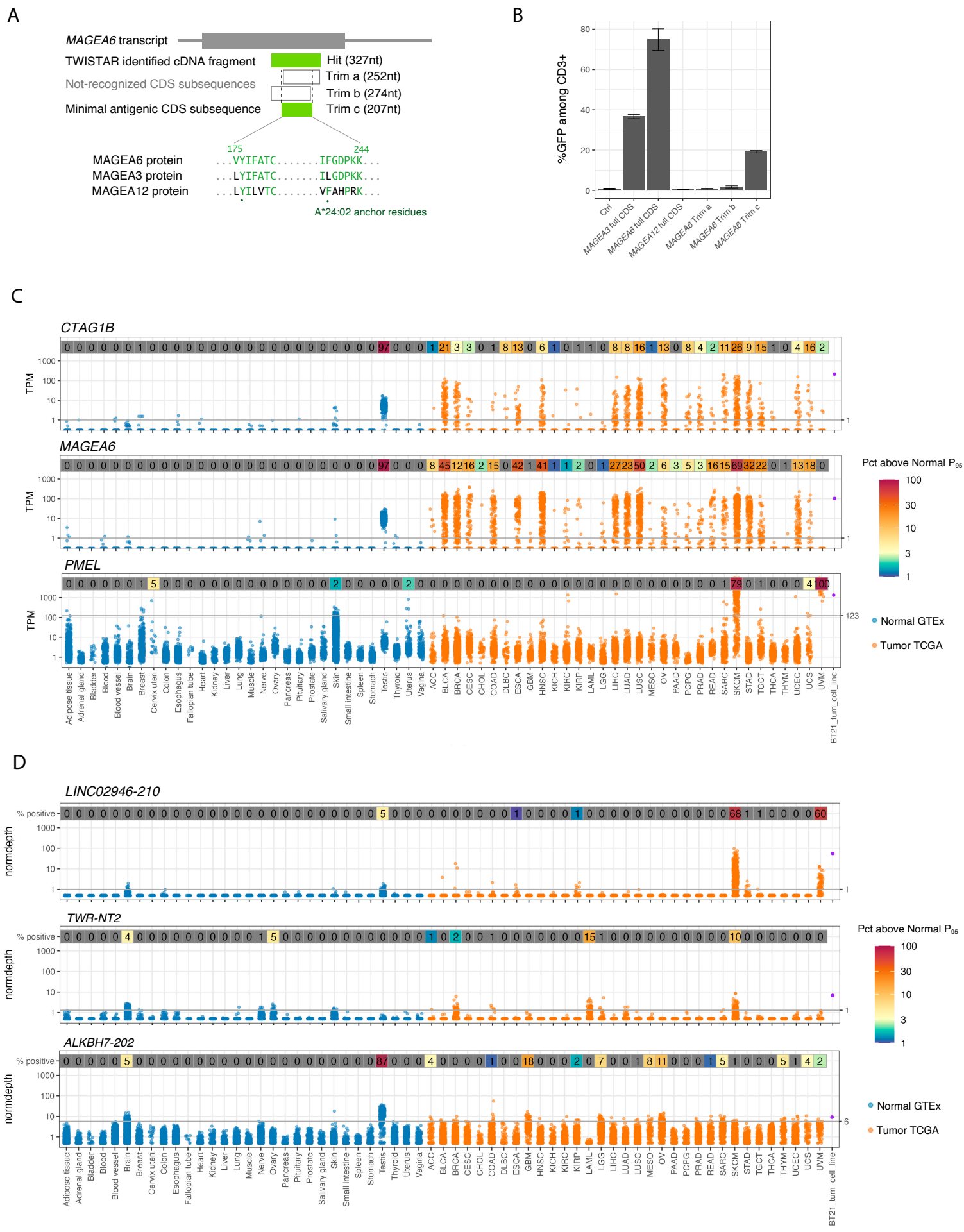

Supplement figure 4

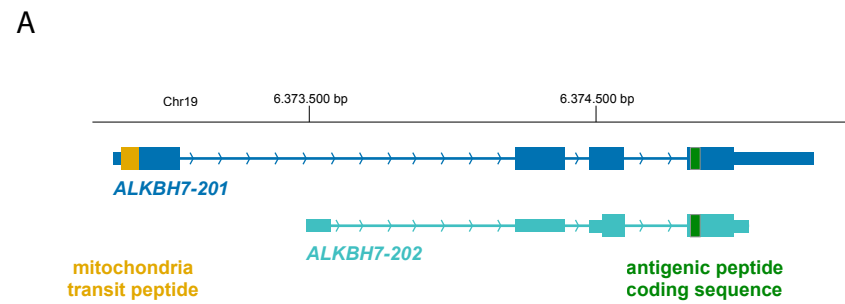

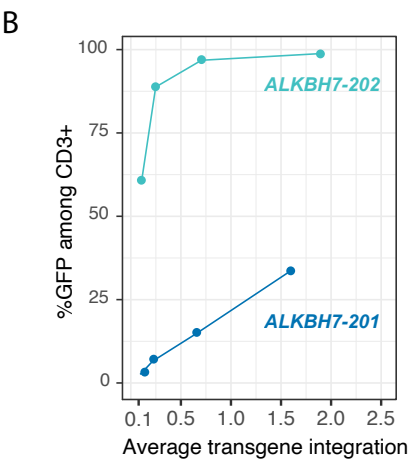

Supplement figure 5

A

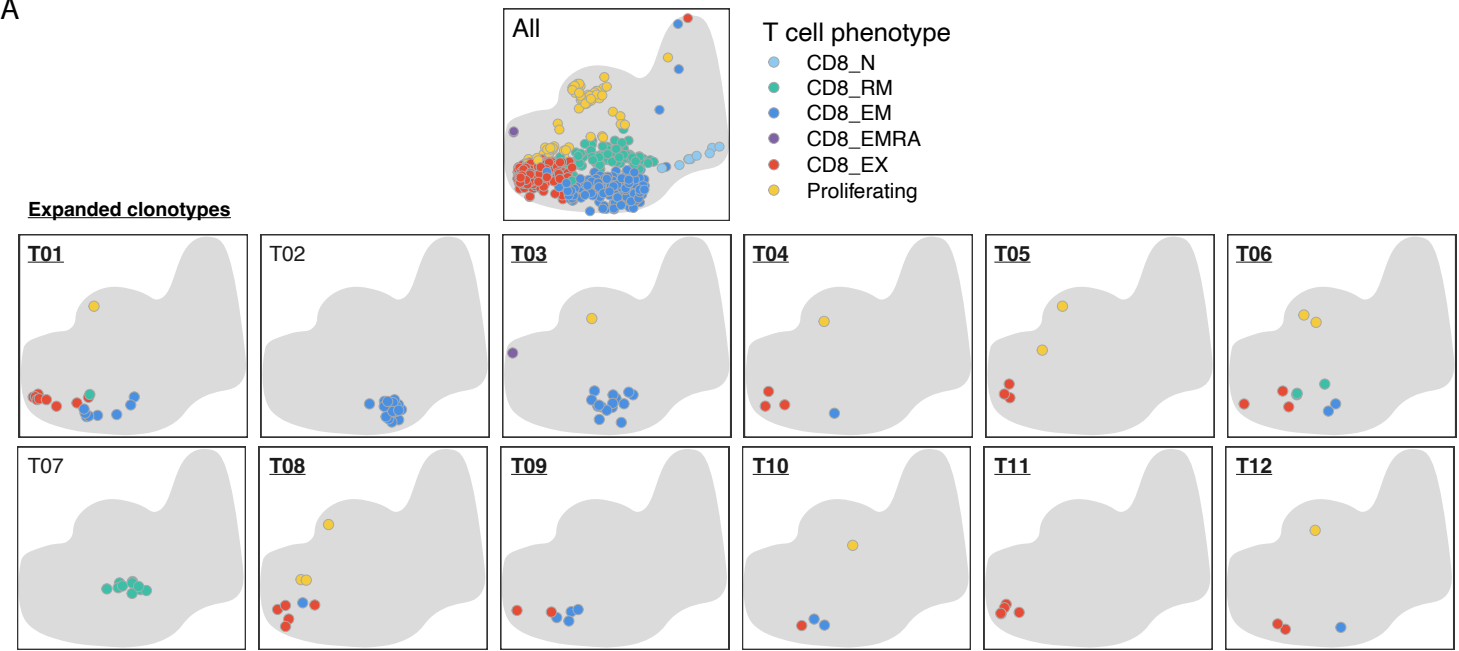

B

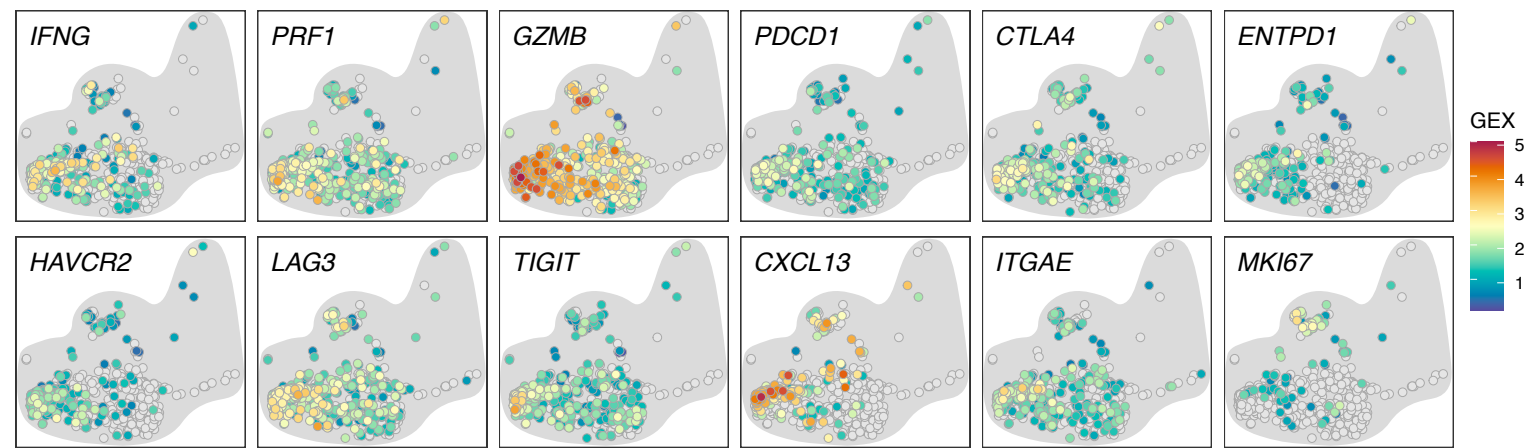

C

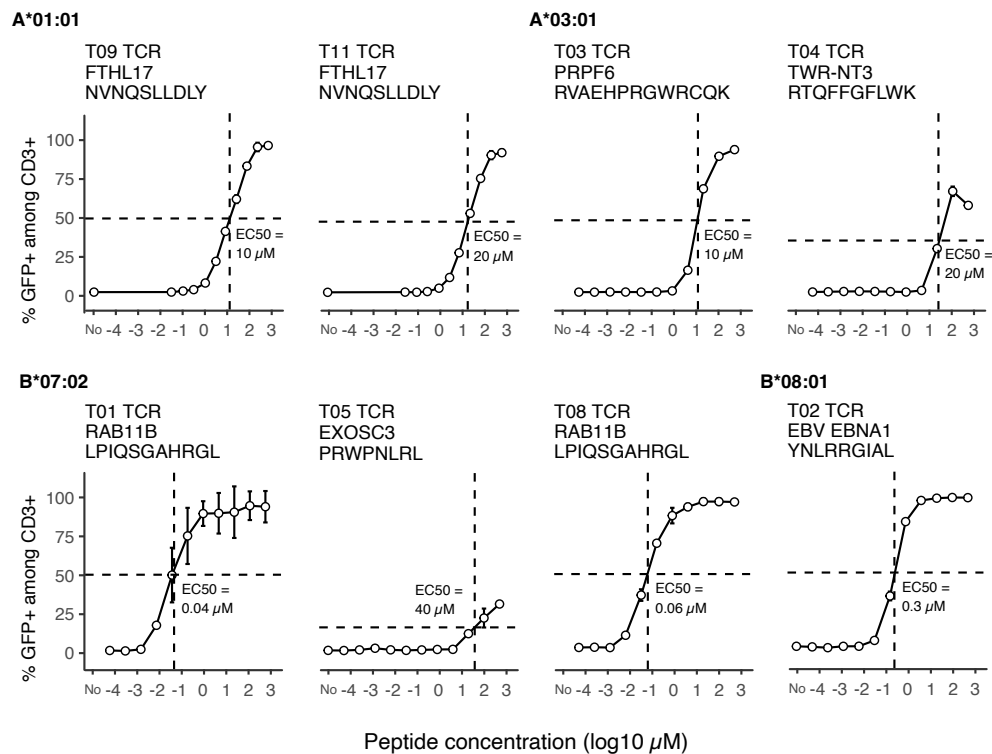

D

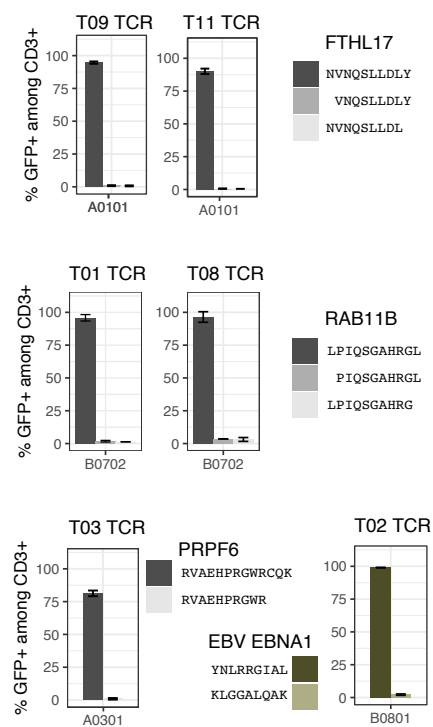

Supplement figure 6

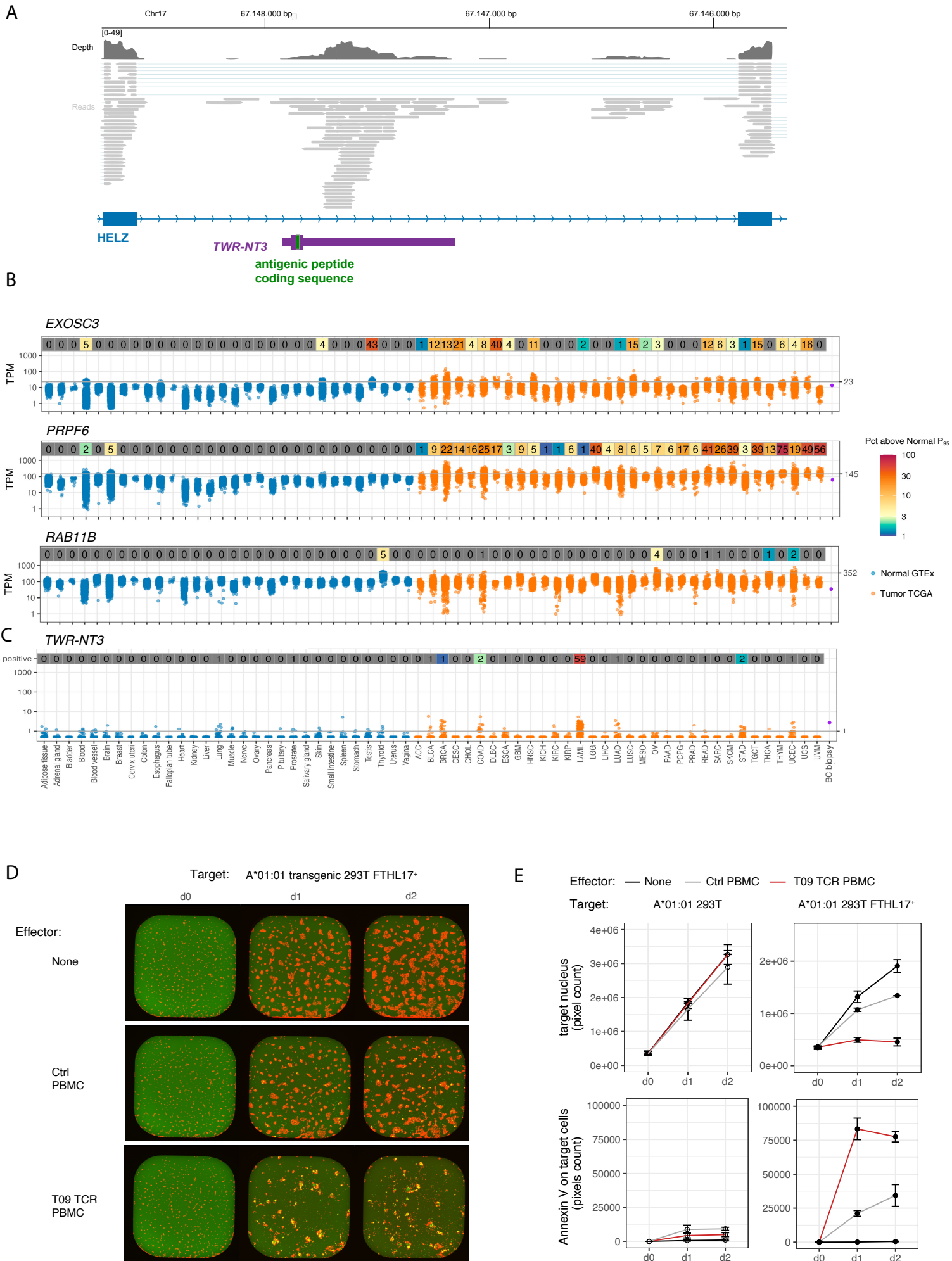

Supplement figure 7

A

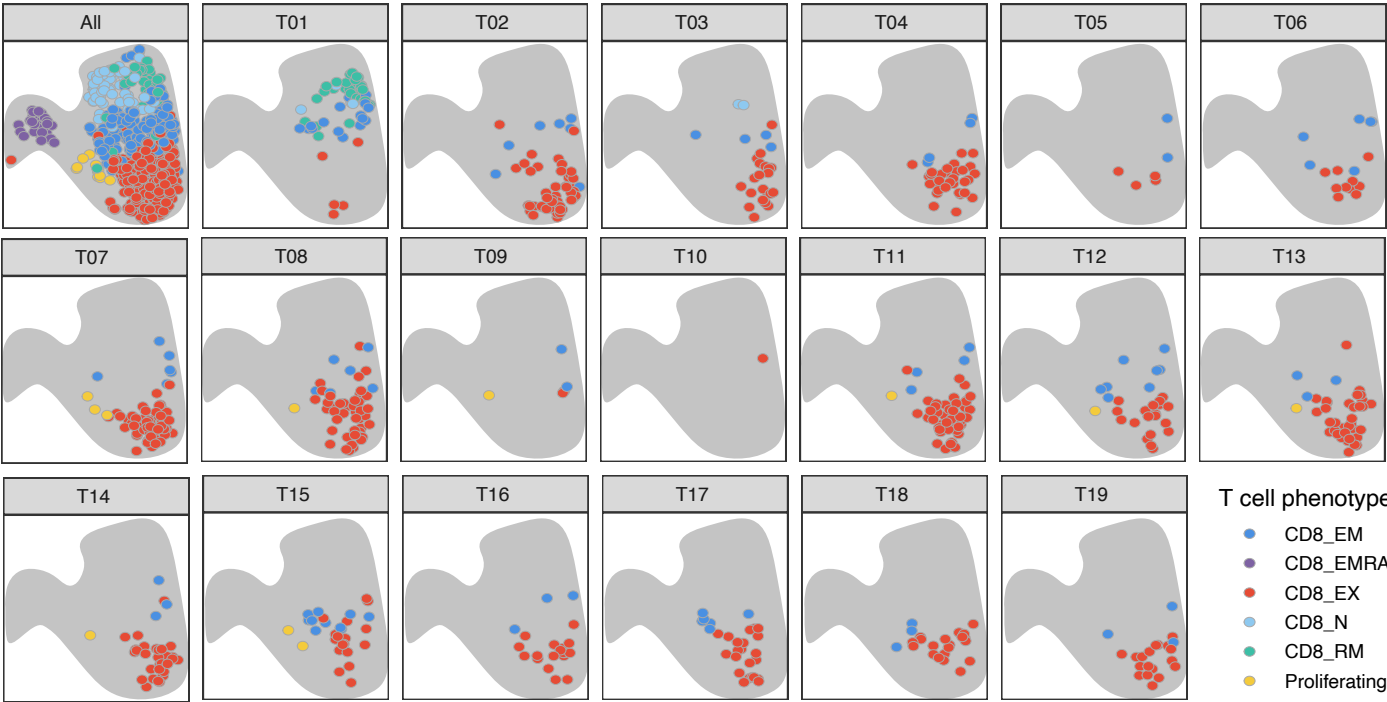

B

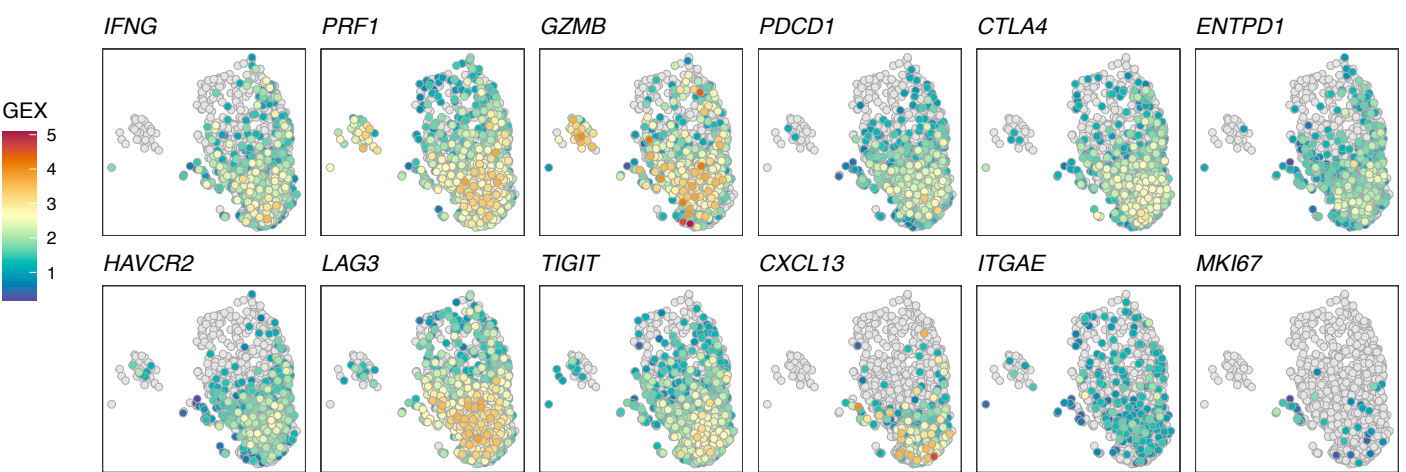

C

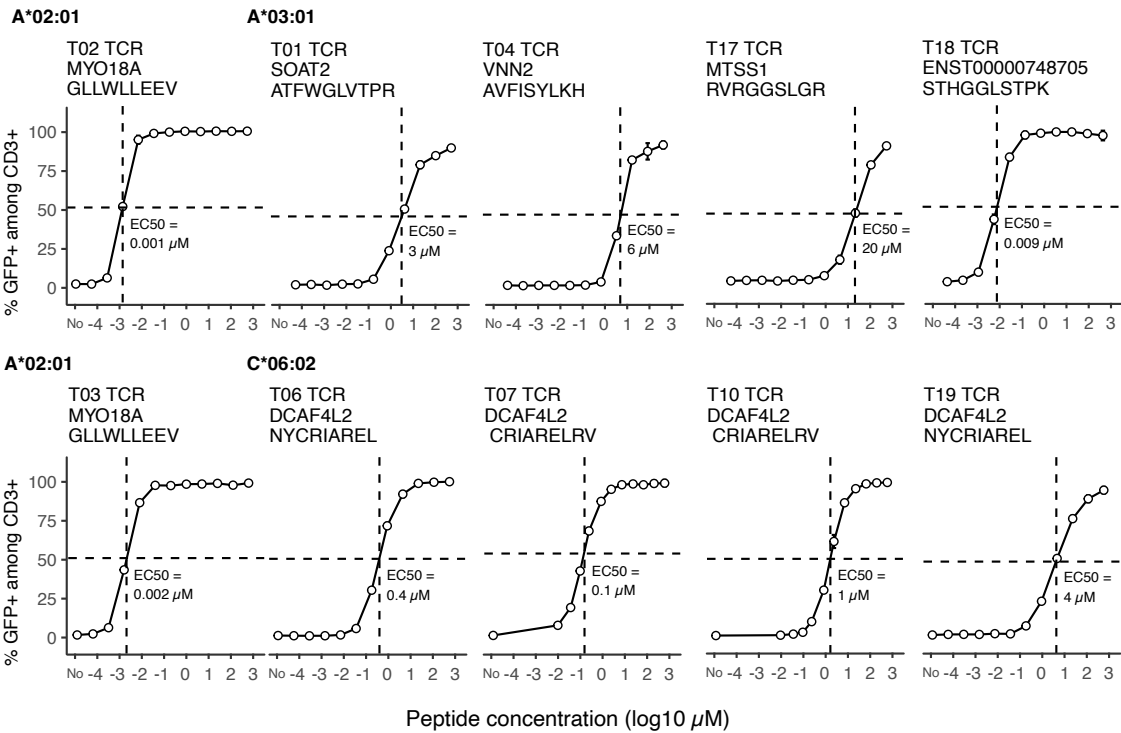

D

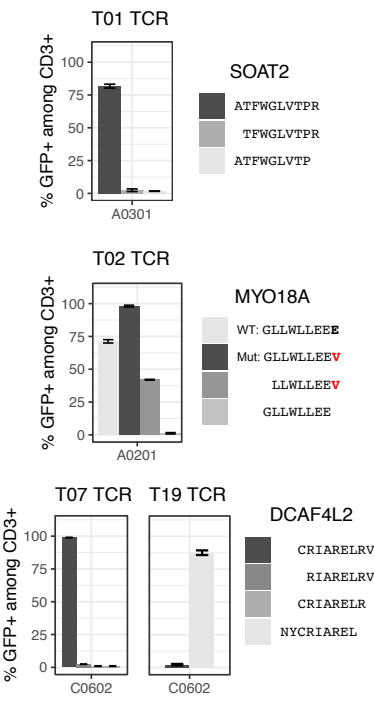

Supplement figure 8

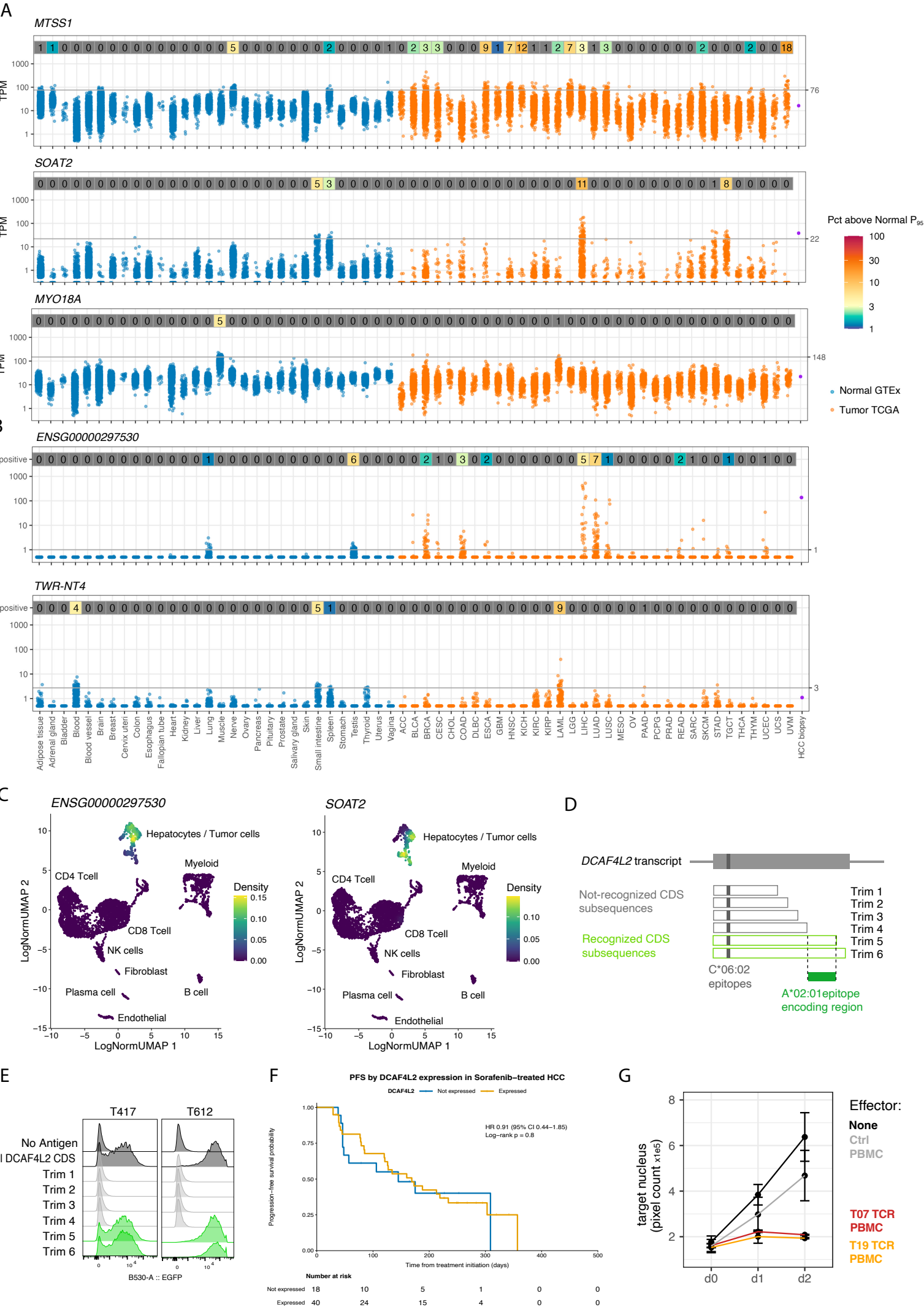
